# Evolutionary Analysis of Gene-expression Localization in the Model Crustacean, *Daphnia pulex*

**DOI:** 10.1101/2025.05.01.651673

**Authors:** Zhiqiang Ye, Tia Swenty, Andrew C. Zelhof, Michael Lynch

## Abstract

Whole-genome sequencing provides lists of genes of putative relevance to organismal biology. However, in all metazoans, a large fraction of inferred genes has no known functions, in some cases with no orthologs in related species, and even orthology at the DNA-sequence level often not providing indisputable evidence of gene function. A first step towards resolving the functional features of gene encyclopedias in multicellular species is to evaluate the tissues in which individual genes are expressed. Here, we report on assays of expression for the full sets of protein-coding and long-noncoding RNA (lncRNA) genes across eight tissues of the microcrustacean *Daphnia pulex*. We also take advantage of a large database on levels of polymorphism and divergence for each gene to infer various features of selection operating on genes expressed in different tissues, including novel genes restricted to particular *Daphnia* lineages. In addition to generating a resource for future work on the molecular, cellular, and developmental biology of the model species *D. pulex*, this study highlights a number of novel findings. These include the identification of sets of genes experiencing unusual forms of positive selection, the discovery of unusual patterns of evolution in the pool of testes-specific genes, rapid turnover and sequence evolution of lncRNA genes, and the pervasive operation of selection on genes thought to be *D. pulex*-specific.

## Introduction

The era of comparative genome and transcriptome sequencing has enabled the identification and provisional annotation of the genes in a large number of species. However, a substantial fraction of genes in most genomes are lineage-specific and/or have undescribed functions, and even those with clear orthologs in different species may not always have shared functions (Gabaldon and Koonin 2013; Weisman et al. 2022). For metazoans, a first step toward improving this situation is to evaluate the locations of gene expression at the tissue level. Without further information at the molecular/cellular levels, this does not reveal the precise structural and/or functional attributes of gene products, but it does narrow down the possibilities. Further insights can be gained when population-genomic data are available for a species, as this can provide information on the strength and patterns of selection operating on various gene products, and comparative analyses across species can yield insight into the stability of gene residence and function over evolutionary time (Lynch 2007).

Here, we examine the tissue-specific expression of the full set of protein-coding genes and noncoding RNAs in the microcrustacean *Daphnia pulex,* in an effort to further establish this species as a central model system in animal biology. A useful feature of this species is the presence of a very large database on within-population variation (nearly 2000 sequenced genomes) and between-species divergence across the entire genome, including information on the presence of lineage-specific genes (Maruki et al. 2022; Ye et al. 2023; Lynch et al. 2024).

Together, these data provide the basis for an encyclopedia of tissue-specific locations of gene expression and the relationship of such localization to levels of polymorphism and divergence at the DNA level. For protein-coding genes, comparison of the behavior of synonymous and nonsynonymous sites yields insight into the degree of selection operating at the amino-acid sequence level. For noncoding RNAs, which comprise a large fraction of the *Daphnia* gene set but whose functions remain largely unknown in any multicellular eukaryote, similar insights can be gained by contrasting levels of variation with those for the silent-sites of protein-coding genes (under the assumption that the latter are behaving in a nearly neutral fashion). In addition, we reveal the general relationship between gene-expression levels and rates of molecular evolution, an issue of long-standing interest (Drummond and Wilke 2008; Zhang and Yang 2015), as well as the unique features of tissue-specific and *D. pulex*-specific genes.

## Results

We generated transcriptome data for eight tissues: swimming antenna, brain, carapace, eye, gut, heart, male sensory antennule (at the base of the head), and testes, as well as for sperm. All samples were collected from adult individuals of *Daphnia pulex* clone KAP4, which also serves as the reference genome (see Methods). For each tissue, we assayed three independent replicates, each generated from 100 individuals. Dissections, RNA extractions, and sequencing were conducted separately to ensure accuracy and reproducibility, with replicate correlations exceeding 0.92. After filtering out unmapped reads, reads with multiple genomic matches, and reads mapped to ribosomal RNA, we obtained an average of 26 million uniquely mapped reads (156× average coverage) for each sample (**Supplementary Table S1**). For consistency and ease of analysis, we use the average expression value from the three biological replicates to represent the gene-expression level in each tissue. Using a minimum cutoff of one transcript per million reads (TPM=1) for each sample, 13,778 (90%) of the annotated 15,282 protein-coding genes for KAP4, were found to be expressed in at least one tissue. Similarly, using a cutoff of TPM = 0.2, 3692 (89%) of the annotated 4,132 long non-coding RNAs (lncRNAs) were expressed in at least one tissue. In the following results, we sequentially present summaries on protein-coding genes and lncRNAs.

### Protein-coding Genes

Among tissues, testes exhibited the lowest number of expressed protein-coding genes (9,411), whereas the male antennule showed the highest (11,408) (**Figure 1**). In total, 7,126 genes were expressed across all tissues, whereas each tissue displayed between 40 and 304 tissue-specific genes. Of the genes expressed within each tissue, the highest incidences of tissue-specific protein-coding genes were found in gut (3.0%) and testes (2.8%). Of the total pool of 1,227 tissue-specific genes, 24.8% were expressed in gut, 21.1% in testes, 14.2% in male antenna, 10.3% in second antenna, 9.3% in brain, 4.6% in carapace, 3.3% in eye, and 2.5% in heart. In general, the numbers of tissue-specific genes with orthologs in the close outgroup *D. obtusa* are 1.1 to 2.8× higher than those without orthologs, but the situation is reversed for testes-specific genes, as the numbers without orthologs are inflated by 30% relative to those with outgroup orthologs (**Table 1**).

**Figure 1.**
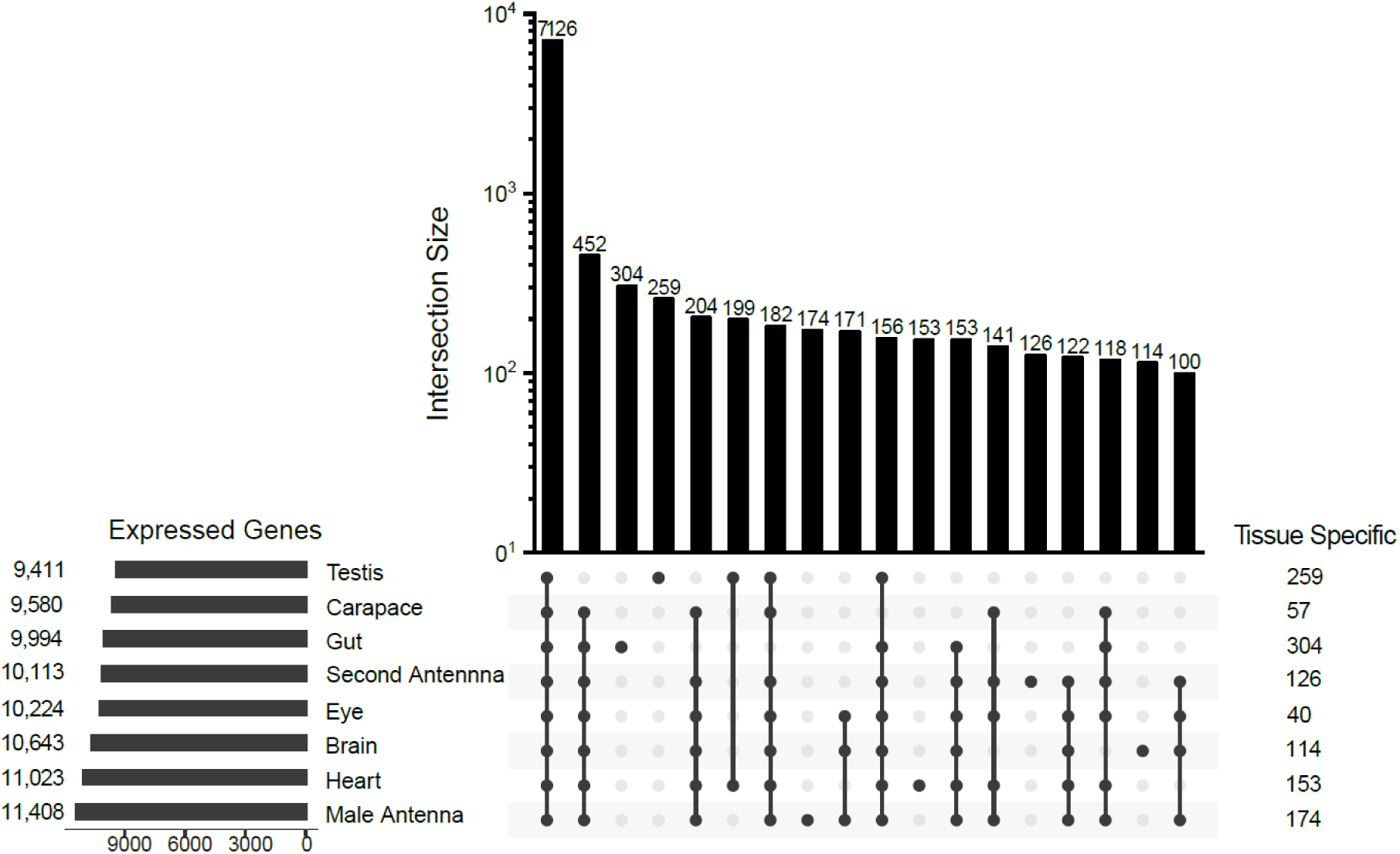
UpSet plot (Lex et al. 2014; Conway et al. 2017) showing the number of unique and shared transcribed protein-coding genes in each tissue, with a minimum cutoff of one transcript per million reads (TPM>1) being used as a criterion for biologically significant expression. Only the twenty most common groupings are shown.

**Table 1.** Average population-genetic parameter estimates associated with protein-coding genes expressed in different tissues. TPM denotes transcripts per million. The top half of the table gives results for the total pool of genes expressed in a tissue; bottom half refers to subsets of genes with tissue-specific expression. Genes are further subdivided into categories with and without orthologous copies in the genome of the close relative *D. obtusa*. Numbers in parentheses in the Genes column refer to the numbers of genes with π_N_ /π_S_ > 10.0, which were not included in the averages reported for this parameter.

| | Genes | TPM | $\pi_N$ | $\pi_S$ | $\pi_N/\pi_S$ | Genes | TPM | $\pi_N$ | $\pi_S$ | $\pi_N/\pi_S$ |
| --- | --- | --- | --- | --- | --- | --- | --- | --- | --- | --- |
|  | General, with orthologs: |  |  |  |  | General, without orthologs: |  |  |  |  |
| Swim. antenna | 8826 (19) | 84.3 | 0.0020 | 0.0158 | 0.208 (0.002) | 1287 (36) | 161.4 | 0.0047 | 0.0140 | 0.875(0.041) |
| Brain | 9202 (22) | 88.9 | 0.0021 | 0.0157 | 0.201 (0.002) | 1441 (43) | 96.1 | 0.0048 | 0.0131 | 0.912(0.038) |
| Carapace | 8355 (19) | 80.8 | 0.0021 | 0.0155 | 0.213 (0.002) | 1225 (37) | 223.5 | 0.0046 | 0.0134 | 0.885(0.042) |
| Eye | 8943 (21) | 62.4 | 0.0020 | 0.0157 | 0.204 (0.002) | 1281 (39) | 321.5 | 0.0047 | 0.0135 | 0.882(0.039) |
| Gut | 8612 (17) | 94.0 | 0.0022 | 0.0160 | 0.212 (0.002) | 1382 (35) | 83.9 | 0.0049 | 0.0145 | 0.836(0.037) |
| Heart | 9417 (20) | 68.3 | 0.0022 | 0.0160 | 0.217 (0.002) | 1606 (38) | 191.4 | 0.0050 | 0.0142 | 0.832(0.034) |
| Male antenna | 9850 (22) | 75.5 | 0.0022 | 0.0164 | 0.211 (0.002) | 1558 (39) | 122.3 | 0.0050 | 0.0142 | 0.877(0.036) |
| Sperm | 9160 (18) | 61.2 | 0.0023 | 0.0160 | 0.220 (0.002) | 1653 (36) | 238.3 | 0.0052 | 0.0136 | 0.856(0.034) |
| Testes | 8034 (16) | 67.7 | 0.0021 | 0.0150 | 0.219 (0.002) | 1377 (32) | 302.5 | 0.0051 | 0.0129 | 0.861(0.035) |
| Mean | 8933 | 75.9 | 0.0021 | 0.0158 | 0.213 | 1423 | 193.4 | 0.0049 | 0.0137 | 0.868 |
| SE | 185 | 3.9 | 0.0000 | 0.0001 | 0.002 | 51 | 28.6 | 0.0001 | 0.0002 | 0.008 |
|  | Specific, with orthologs: |  |  |  |  | Specific, without orthologs: |  |  |  |  |
| Swim. antenna | 80 (0) | 26.2 | 0.0026 | 0.0260 | 0.222 (0.024) | 46 (0) | 32.6 | 0.0029 | 0.0158 | 1.244(0.405) |
| Brain | 62 (1) | 5.2 | 0.0037 | 0.0220 | 0.345 (0.101) | 52 (1) | 2.3 | 0.0025 | 0.0083 | 1.318(0.432) |
| Carapace | 42 (0) | 8.0 | 0.0059 | 0.0277 | 0.231 (0.022) | 15 (0) | 7.4 | 0.0069 | 0.0186 | 0.459(0.108) |
| Eye | 21 (0) | 5.4 | 0.0036 | 0.0178 | 0.292 (0.081) | 19 (0) | 145.9 | 0.0024 | 0.0055 | 0.781(0.258) |
| Gut | 198 (0) | 47.7 | 0.0057 | 0.0255 | 0.299 (0.021) | 106 (0) | 44.1 | 0.0076 | 0.0270 | 0.350(0.031) |
| Heart | 93 (1) | 11.6 | 0.0055 | 0.0258 | 0.323 (0.041) | 60 (0) | 13.8 | 0.0050 | 0.0198 | 0.748(0.186) |
| Male antenna | 125 (1) | 41.0 | 0.0023 | 0.0255 | 0.228 (0.018) | 49 (1) | 10.2 | 0.0052 | 0.0231 | 0.427(0.094) |
| Testes | 113 (0) | 16.0 | 0.0058 | 0.0204 | 0.398 (0.038) | 146 (1) | 17.6 | 0.0075 | 0.0156 | 0.652(0.061) |
| Mean | 92 | 20.1 | 0.0044 | 0.0239 | 0.292 | 62 | 34.3 | 0.0058 | 0.0167 | 0.747 |
| SE | 20 | 5.8 | 0.0005 | 0.0012 | 0.022 | 16 | 16.7 | 0.0005 | 0.0025 | 0.129 |

Aside from their relevance to the biology of the organism, the identities of these tissue-specific genes may be useful in future single-cell RNA-sequencing analyses involving whole organisms that bypass laborious tissue dissections (Krishnan et al. 2024). A limitation of the latter type of study is the need to indirectly infer the ultimate locales of cell-type clusters from the known expression patterns of genes in other (often distantly related) species. Annotated tissue-specific genes offer a way to minimize such problems, by providing markers for the tissue locations for otherwise anonymous individual cells.

### Evolutionary analysis

A useful resource for the *D. pulex* system is the availability of substantial databases on levels of polymorphism and divergence for the full set of protein-coding genes. These are derived from two large surveys, one based on ∼800 isolates distributed over ten populations (Maruki et al. 2022) and the other based on ten years of data from a single representative population, with ∼90 clonal isolates sequenced per year (Ye et al. 2023; Lynch et al. 2024). To minimize the effects of sampling error, the general evolutionary-genetic features of each gene were obtained by averaging the results over both of these studies, which have been shown to be statistically concordant (Ye et al. 2023). (All gene-specific data are compiled in the **Supplementary File**).

Regarding within-species variation, the results reported on here are based on the pool of genes with π_N_/π_S_ < 10.0, where the numerator and denominator denote the levels of nucleotide diversity (π) at nonsynonymous and synonymous sites, respectively. This restriction, which is made to avoid outliers resulting from spuriously low sample estimates of π_S_ and to avoid any potential mapping errors with respect to paralogous genes, reduces the pool of genes examined at the population-genetic level by only 1%. For genes with orthologs in the outgroup species *D. obtusa*, there is very little difference in mean π_N_/π_S_ among tissues, the range being 0.201 to 0.220 (all with SEs close to 0.002) (**Table 1**). For genes without *D. obtusa* orthologs, mean π_N_/π_S_ is greatly elevated, by a factor of ∼4, but again with little variation among tissues, ranging from 0.832 to 0.912 (with average SE = 0.037). Although estimates of π_N_/π_S_ > 10.0 were excluded from these analyses, consistent with the already very high mean π_N_/π_S_ for the pool of genes without *D. obtusa* orthologs, there is a substantial and systematic difference in the fraction of such outlier genes between the pools with and without *D. obtusa* orthologs, 0.2 and 2.6% respectively (**Table 1**).

The small sets of tissue-specific genes have different evolutionary features than the bulk pool of genes. For those with *D. obtusa* orthologs, π_N_ is inflated by an average factor of 2.3×, relative to that for genes with more general expression, whereas π_S_ is inflated by 1.5×, leading to an average inflation of π_N_/π_S_ of ≃ 1.4× (**Table 1**). For tissue-specific genes without *D. obtusa* orthologs, the SEs for π_N_/π_S_ are larger owing to the smaller numbers of genes, and such a distinction cannot be made.

Based on levels of sequence divergence from *D. obtusa* orthologs, the average features of the total pools of expressed genes in each tissue are similar (**Table 2**). Mean estimates of within-tissue *d*_N_ fall in the range of 0.017 to 0.019, whereas those for *d*_S_ are all ≃ 0.13, and all estimates of *d*_N_/*d*_S_ fall in the range of 0.13 to 0.14. For the tissue-specific genes, there is slight but consistent elevated divergence, with mean estimates of *d*_N_ in the range of 0.024 to 0.036, except for testes (where *d*_N_ ≃ 0.047), whereas *d*_S_ falls in the range of 0.15 to 0.17, and *d*_N_/*d*_S_ in the range of 0.16 to 0.26 (except for testes, where the ratio is substantially higher, at ≃ 0.33).

**Table 2.** Average sequence divergence estimates for protein-coding genes with orthologous copies in *Daphnia* obtusa. For genes with expression in two or more tissues, the SEs of all *d*_N_ and *d*_S_ are 0.0003 and 0.0005, respectively. For genes with tissue-specific expression, the SEs of the tissue-specific mean *d*_N_ range from 0.0012 to 0.0040, and for *d*_S_ range from 0.0035 to 0.0176. Note that the tissue-specific data for sperm are essentially the same as those for testes.

| | $d_N$ | $d_S$ | $d_N/d_S$ (SE) | $d_N$ | $d_S$ | $d_N/d_S$ (SE) |
| --- | --- | --- | --- | --- | --- | --- |
|  | General: |  |  | Tissue-specific: |  |  |
| Swim. antenna | 0.0176 | 0.1273 | 0.129 (0.002) | 0.0288 | 0.1678 | 0.190 (0.017) |
| Brain | 0.0178 | 0.1270 | 0.132 (0.002) | 0.0277 | 0.1491 | 0.197 (0.026) |
| Carapace | 0.0183 | 0.1273 | 0.133 (0.002) | 0.0372 | 0.1555 | 0.257 (0.025) |
| Eye | 0.0173 | 0.1267 | 0.128 (0.002) | 0.0243 | 0.1609 | 0.161 (0.017) |
| Gut | 0.0184 | 0.1285 | 0.134 (0.002) | 0.0356 | 0.1617 | 0.225 (0.012) |
| Heart | 0.0191 | 0.1287 | 0.140 (0.002) | 0.0345 | 0.1628 | 0.238 (0.018) |
| Male antenna | 0.0184 | 0.1292 | 0.134 (0.002) | 0.0277 | 0.1597 | 0.189 (0.011) |
| Sperm | 0.0194 | 0.1285 | 0.142 (0.002) | ----- | ----- | ----- |
| Testes | 0.0187 | 0.1259 | 0.138 (0.002) | 0.0470 | 0.1612 | 0.334 (0.020) |
| Mean | 0.0183 | 0.1277 | 0.134 | 0.0328 | 0.1598 | 0.224 |
| SE | 0.0002 | 0.0004 | 0.002 | 0.0026 | 0.0020 | 0.019 |

Even in the best-characterized metazoan genomes, many predicted genes inferred to be active based on expression evidence are nonetheless unannotated, because they lack obvious orthologs in well-characterized model species (e.g., determined by applying Blast to all protein-coding gene models within NCBI, as done here). For the 4,085 unannotated protein-coding genes in the KAP4 genome (inferred to exist based on the presence of open reading frames and prior expression data from whole individuals), 3,319 were expressed in at least one tissue, whereas 766 had insignificant expression. In these two respective groups with population-genomic data, average π_N_/π_S_ is 0.396 (0.014; N = 2,835) and 0.736 (0.059; N = 426), far higher than observed for the pool of protein-coding genes with *D. obtusa* orthologs but lower than that for the general group without orthologs (**Table 1**). Both groups have average *d*_N_/*d*_S_ estimates that are about double those observed for annotated genes (**Table 2**), 0.235 (0.004) and 0.252 (0.014) respectively, and *d*_S_ is slightly elevated as well, 0.138 (0.001) and 0.195 (0.007) respectively. Taken together, the fact that average levels of π_N_/π_S_ and *d*_N_/*d*_S_ are both < 1 suggests that the bulk of unannotated genes in KAP4 are either currently experiencing selection, or were so early in the history of the *D. pulex* lineage.

Of the pool of unannotated protein-coding genes with expression data (**Supplementary Table S2**), the 2,385 with orthologs in *D. obtusa* have mean π_N_/π_S_ = 0.304 (0.009), whereas the 450 without orthologs have π_N_/π_S_ = 0.776 (0.046), suggesting somewhat relaxed purifying selection in the latter group. Of the pool of unannotated genes with no evidence of expression, the 268 genes with *D. obtusa* orthologs have mean π_N_/π_S_ = 0.549 (0.057), whereas the 158 without orthologs have π_N_/π_S_ = 1.054 (0.121). Thus, taken as a group, unannotated genes lacking in expression are experiencing weaker levels of purifying selection, and the small subset also lacking *D. obtusa* orthologs are pseudogene candidates. To be more precise, letting joint values of π_N_/π_S_ and *d*_N_/*d*_S_ that are not significantly different from 1.0 be the hallmark of a neutrally evolving gene, for the pool of expressed unannotated genes, just 206 (7.3%) meet the criteria of being pseudogenes, and this increases to 84 (19.7%) for the unexpressed unannotated genes.

In an effort to characterize genes with strong evidence for unusual patterns of positive selection, we focused on the subset of 24 genes with *d*_N_/*d*_S_ significantly greater than 1.0 (**Supplementary Table S3**). Notably, most of these genes also have very high π_N_/π_S_, often significantly greater than the neutral expectation of 1.0. Twelve of them are uncharacterized, in that they are without orthologs with known functions in other model metazoan species. The most rapidly diverging protein by far (LOC124194853), with *d*_N_/*d*_S_ = 4.99 (0.39), is annotated as “DNA-directed RNA polymerase II subunit RPB1-like,” which implies involvement in transcription of protein-coding and/or lncRNA genes. Inferred based on data in the UniProt database, this gene is not restricted to *D. pulex*, as it is found in other *Daphnia* species. It is expressed in all tissues (but ∼ 3× more in testes and sperm), and also has an exceptionally high estimate of π_N_/π_S_ = 2.26 (0.29), although this is substantially lower than the divergence ratio, which implies positive selection. The second most highly diverging protein-coding gene, with *d*_N_/*d*_S_ = 2.36 (0.07) and π_N_/π_S_ = 1.23 (0.04) again significantly elevated above the neutral expectation, is a sucrase-isomaltase with putative functions of breaking down sucrose and maltose into simpler sugars; it too is ubiquitously expressed, but with ∼60-fold expression elevation in the gut. In the above cases, both π_N_/π_S_ and *d*_N_/*d*_S_ significantly exceed the neutral expectation of 1.0, suggesting selection for both divergence among species and maintenance of diversity within species at the amino-acid level.

Focusing now on outliers with respect to within-species diversity, 214 genes meet the criterion of having π_N_/π_S_ estimates significantly >1.0 (**Supplementary File**). Of these, 126 (59%) have no orthologs in *D. obtusa*, well above the 14% observed across the entire genome. Two of these genes have already been noted above. We have pointed out before that outlier genes in *D. pulex* with high π_N_/π_S_ have diverse functions, but are enriched with those involved in DNA/RNA processing, mitochondrial function, neurotransmission, development, and lysosome activity (Ye et al. 2023). For example, of the 16 annotated genes with the highest values of π_N_/π_S_ (ranging in decreasing order from 8.53 to 2.90, and all with broad expression unless explicitly noted) that have not already been mentioned, six have diverse functions associated with DNA/RNA metabolism: cohesin subunit SA-1 (mediates sister-chromatid cohesion during mitosis and meiosis; nearly exclusively expressed in testes); glycine-tRNA ligase; tRNA pseudouridine synthase (involved in post-translational modification of tRNAs); helicase/nuclease DNA2 (involved in Okazaki-fragment processing in DNA replication and repair); RNA-binding protein FUS (involved in pre-mRNA splicing; heavily expressed in antennae and carapace); and S-methyl-5’- thioadenosine phosphorylase (involved in the salvage pathways for adenine and methionine). An additional four members of this group of 16 have diverse mitochondrial functions: holocytochrome c-type synthase (catalyzes heme-group attachment to cytochrome c in mitochondria); CYFIP-related Rac1 interactor B (involved in mitochondrial GTPase activity); carnitine O-palmitoyltransferase 2 (involved in mitochondrial fatty-acid metabolism); and NADH-ubiquinone oxidoreductase 75 kDa subunit (couples electron transfer from NADH to ubiquinone in the mitochondrial electron transport chain). This selected list of proteins confirms our prior conclusions as to the functional groupings of genes that are under apparent strong balancing selection within populations.

Finally, we consider whether the evolutionary-genetic features of protein-coding genes are associated with levels of expression. For genes with *D. obtusa* orthologs, within each tissue, there is a weak but significant negative association between all of the above population-genetic parameters and expression level (**Supplementary File**). Using log-transformed data, the average slope for the regression involving π_N_ is -0.25 (with a range of -0.30 to -0.15 among tissues), all with SEs ≃ 0.008, significance values < 10^−11^, and average r^2^ = 0.12. Thus, on average, for every 10^4^-fold increase in expression level, there is a ten-fold decrease in π_N_. The responses are weaker for π_S_, but nonetheless negative, with an average slope for the regressions of -0.10 (with a range of -0.14 to -0.05 among tissues), all with SEs ≃ 0.006, significance values < 10^-11^, and average r^2^ = 0.03. For π_N_/π_S_, the average slope for the regressions is -0.14 (with a range of -0.16 to -0.11 among tissues), all with SEs ≃ 0.008, significance values < 10^−11^, and average r^2^ = 0.03.

These same patterns of dependence on gene expression appeared for levels of sequence divergence from the outgroup species (**Supplementary File**). The slopes and level of correlation are essentially the same for π_N_ and *d*_N_, whereas the slope is ∼2× steeper for π_S_ than for *d*_S_ (with a similar average correlation). However, the average regressions for *d*_N_/*d*_S_ are ∼2-fold steeper than those for π_N_/π_S_ (both negative). Thus, regardless of the measure of divergence or efficiency of selection on amino-acid sequences, all observations point to the average strength of purifying selection being elevated in more highly expressed protein-coding genes.

## Noncoding RNAs

Long non-coding RNAs (lncRNAs) are conventionally defined as RNA molecules >200 nucleotides in length that lack the ability to code for proteins. As for protein-coding genes, they are transcribed by RNA polymerase II, and they have structures similar to messenger RNAs (mRNAs), including the common presence of spliceosomal introns and poly-A tails. The functions of lncRNA molecules are mostly unknown but thought to be involved in diverse aspects of gene-expression regulation, including interactions with DNA, protein, and mRNA, and in chromatin remodeling.

Numerous (4,132) lncRNA genes have been suggested to reside in the KAP4 genome based on previous whole-organism expression data (NCBI accession: PRJNA777597). We found 3,692 such genes to be expressed in at least one tissue, 10% of which have no apparent orthologs in *D. obtusa*. Most such genes were expressed in more than one tissue type, and 916 were found to be expressed in all tissues (**Figure 2, Supplementary File**). The smallest numbers of expressed lncRNAs (∼1,700 to 1,800) were associated with carapace, swimming antennae, and male antennules, whereas the numbers in sperm and testes were substantially elevated (∼2,800 each). The average number of tissue-specific lncRNAs detected per tissue is just 19, dramatically lower than that for protein-coding genes, excepting the very high number in testes (317).

**Figure 2.**
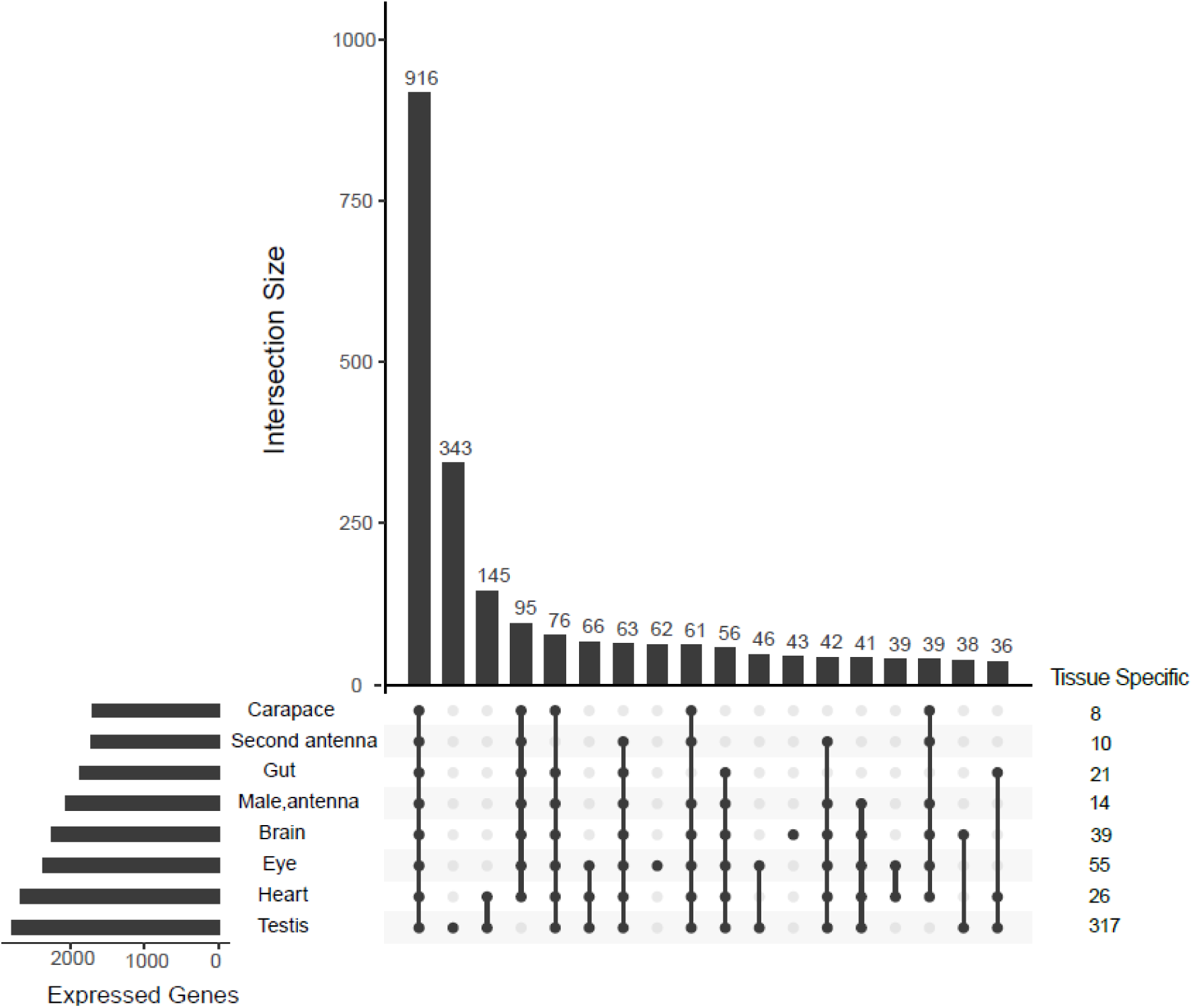
UpSet plot showing the number of unique and shared transcribed long-noncoding RNA genes in each tissue, with a minimum cutoff of TPM>0.2 being used as a criterion for biologically significant expression. Only the eighteen most common groupings are shown.

Among the 4,132 lncRNAs, 1,122 have genomic locations that are overlapping with those for protein-coding genes, with the remainder being located in intergenic regions. Of the former, 1,007 are antisense lncRNAs and 115 are sense-overlapping lncRNAs. 785 of the lncRNAs overlap with introns in protein-coding genes, although none are entirely embedded within introns.

### Evolutionary analysis

The preceding survey indicates that lncRNAs are prominent contributors to the *D. pulex* transcriptome, and here we use population-genomic data to obtain insight into the strength of selection operating on such enigmatic sequences. Population-level sequence analysis indicates that on average lncRNAs have substantially less average nucleotide diversity (π = 0.0097 and 0.0080 for genes with and without *D. obtusa* orthologs, with little variation among tissues; **Table 3**) than levels of silent-site diversity in protein-coding genes (0.0158 and 0.0137, respectively, from **Table 1**). Average π for lncRNAs is somewhat higher in the tissue-specific noncoding RNAs (0.0119 for those with *D. obtusa* orthologs), but still lower than that seen for π_S_ in protein-coding genes. The average level of nucleotide divergence (contrasted against *D. obtusa*) falls consistently in the narrow range of 0.057 to 0.060 for generally expressed lncRNA genes. This is greater than the level observed for amino-acid replacement sites, which falls in the range of 0.017 to 0.019 for broadly expressed protein-coding genes, but less than that for silent-site divergence for such protein-coding genes, which consistently falls in the range of 0.126 to 0.128 (**Table 2**). For lncRNA genes with tissue-specific expression, the level of divergence falls in the range of 0.065 to 0.077, again higher than that for *d*_N_ for such genes (range 0.024 to 0.034) but lower than that for *d*_S_ (range 0.149 to 0.168).

**Table 3.** Average population-genetic parameter estimates associated with long noncoding RNA genes expressed in different tissues. TPM denotes the mean number/million of transcripts per expressed gene, normalized to a transcript of 1-kb length. The top half of the table gives results for the total pool of genes expressed in a tissue; bottom half refers to subsets of genes with tissue-specific expression. Genes are further subdivided into categories with and without orthologous copies in the genome of the close relative *D. obtusa*. Standard errors are in parentheses. TPM = 0.2 was taken as an indicator of lncRNA gene expression.

| | Genes | TPM | $\pi$ | $d$ | Genes | TPM | $\pi$ |
| --- | --- | --- | --- | --- | --- | --- | --- |
| General, with orthologs: |  |  |  |  | General, without orthologs: |  |  |
| Swim antenna | 1117 | 35.4 (6.8) | 0.0096 (0.0002) | 0.057 (0.001) | 584 | 11.0 (3.9) | 0.0080 (0.0002) |
| Swim antenna | 1117 | 35.4 (6.8) | 0.0096 (0.0002) | 0.057 (0.001) | 584 | 11.0 (3.9) | 0.0080 (0.0002) |
| Brain | 1352 | 23.3 (4.1) | 0.0095 (0.0001) | 0.058 (0.001) | 897 | 10.1 (0.7) | 0.0079 (0.0002) |
| Carapace | 1056 | 40.7 (10.0) | 0.0097 (0.0002) | 0.058 (0.001) | 630 | 7.8 (1.8) | 0.0080 (0.0002) |
| Eye | 1489 | 15.4 (4.2) | 0.0096 (0.0001) | 0.058 (0.001) | 874 | 3.8 (0.4) | 0.0080 (0.0002) |
| Gut | 1565 | 40.6 (22.4) | 0.0097 (0.0001) | 0.059 (0.001) | 703 | 10.8 (6.5) | 0.0081 (0.0002) |
| Heart | 1322 | 30.1 (6.3) | 0.0097 (0.0001) | 0.059 (0.001) | 1090 | 5.8 (1.0) | 0.0080 (0.0001) |
| Male antenna | 1117 | 45.4 (11.4) | 0.0096 (0.0002) | 0.058 (0.001) | 728 | 11.0 (3.2) | 0.0079 (0.0002) |
| Sperm | 1470 | 14.7 (4.2) | 0.0101 (0.0001) | 0.060 (0.001) | 1308 | 11.4 (2.8) | 0.0081 (0.0001) |
| Testes | 1594 | 16.4 (3.1) | 0.0101 (0.0001) | 0.060 (0.001) | 1353 | 12.4 (2.0) | 0.0081 (0.0001) |
| Mean | 1342 | 29.1 | 0.0097 | 0.0585 | 907 | 9.3 | 0.0080 |
| SE | 68 | 4.0 | 0.0000 | 0.0003 | 95 | 1.0 | 0.0000 |
| Specific, with orthologs: |  |  |  |  | Specific, without orthologs: |  |  |
| Swim antenna | 6 | 4.0 (2.9) | 0.0111 (0.0018) | 0.073 (0.006) | 4 | 6.82 (6.28) | 0.0053 (0.0011) |
| Brain | 7 | 1.7 (0.6) | 0.0120 (0.0016) | 0.076 (0.008) | 32 | 1.57 (0.45) | 0.0085 (0.0013) |
| Carapace | 5 | 2.4 (1.6) | 0.0137 (0.0028) | 0.065 (0.014) | 3 | 3.13 (1.32) | 0.0086 (0.0020) |
| Eye | 28 | 0.6 (0.2) | 0.0089 (0.0010) | 0.072 (0.006) | 27 | 0.44 (0.06) | 0.0076 (0.0008) |
| Gut | 15 | 15.7 (12.3) | 0.0133 (0.0015) | 0.071 (0.006) | 6 | 2.13 (0.80) | 0.0062 (0.0025) |
| Heart | 5 | 1.0 (0.4) | 0.0076 (0.0020) | 0.077 (0.008) | 21 | 0.51 (0.07) | 0.0103 (0.0013) |
| Male antenna | 8 | 19.5 (7.4) | 0.0172 (0.0024) | 0.072 (0.006) | 6 | 1.49 (0.26) | 0.0085 (0.0022) |
| Testes | 68 | 3.2 (0.7) | 0.0113 (0.0007) | 0.070 (0.003) | 249 | 3.99 (0.38) | 0.0081 (0.0003) |
| Mean | 18 | 6.0 | 0.0119 | 0.0722 | 44 | 2.5 | 0.0079 |
| SE | 7 | 2.4 | 0.0010 | 0.0013 | 28 | 0.7 | 0.0005 |

As with protein-coding genes, we find prevalent negative associations between the expression levels of lncRNA genes and gene-specific measures of within-population population-genetic diversity (**Supplementary File**). For 14 of 16 evaluations of tissue-specific gene expression (for genes both with and without *D. obtusa* orthologs), the correlations of expression level and π are negative, although the absolute values of the average slopes and r^2^ values are substantially smaller than those for protein-coding genes (for both π_N_ and π_S_). In contrast, unlike the situation for protein-coding genes, for lncRNA genes with *D. obtusa* orthologs, there is little evidence for correlations between expression level and divergence at the sequence level, with the average regression being not significantly different from zero.

Given that a large fraction of lncRNAs (1,122 of 4,132) overlap at least partially with exonic sequences of protein-coding genes, it is not surprising that they have levels of π and *d* intermediate to estimates for silent and replacement sites in protein-coding genes. However, those with no overlap with protein-coding genes have lower total π than those with overlap, 0.0083 (0.0001) vs. 0.0099 (0.0002), and the respective values for total divergence are very similar, 0.0629 (0.0007) vs. 0.0557 (0.0008). In total, there are 482 π outlier lncRNA genes (defined as having significantly larger π than average π_S_ in protein-coding genes; **Supplementary File**), of which 306 do not overlap with any existing protein-coding genes. However, no *d*-outlier genes were found, possibly because the lncRNAs are much shorter than the protein-coding genes, which results in a reduction in statistical power.

### Two vignettes

The data presented above begin to shed light on organism-wide localization of gene expression and the degree to which tissue-specificity exists within *D. pulex*. Combined with the detailed information that exists for each gene with respect to nucleotide polymorphism and divergence in this species, this then starts to provide a template for understanding the evolutionary features of gene expression and diversification. As the number of areas in which this knowledge base can be applied is very large, here we simply point out two exemplar areas of application.

#### Testes-specific genes

One of the more striking patterns emerging from this analysis is the unique behavior of genes expressed in testes/sperm. Although the latter express smaller numbers of protein-coding genes than do cells in other tissues, the incidence of tissue-specific protein-coding genes is elevated in testes. The latter have elevated rates of nonsynonymous substitution, as well as elevated *d*_N_/*d*_S_ ratios, and an exceptionally high fraction of such genes appears to be absent from the genome of the close relative *D. obtusa*. In addition, there is a two-fold increase in the numbers of expressed lncRNA genes in testes relative to other tissues, and the numbers of tissue-specific lncRNAs in testes is elevated ∼17-fold relative to the situation in other tissues, 80% of which do not have obvious orthologs in *D. obtusa*. These results indicate that testes provide a hotbed of evolutionary activity, enriched for both protein-coding and lncRNA genes that are rapidly evolving, tissue-specific, and lineage-specific.

The mechanisms driving these unusual patterns of evolution remain unclear, although the possibilities include sperm competition and aspects of female choice. It should be kept in mind, however, that most generations (typically >75%) in *D. pulex* consist entirely of parthenogenetic females, during which times testes-specific genes will be free from the operation of selection, which may facilitate the relatively rapid rate of evolution of testes-specific genes. Annual bouts of sexual reproduction (with males being produced by environmental sex determination) almost always occur in *D. pulex*, which inhabits temporary ponds. However, in such generations, the sex ratio is generally still biased towards females, which may reduce male competition for access to females.

Finally, it has been argued that elevated rates of evolution can arise in testes-specific genes if the effective population sizes of male-specific genes are reduced relative to that for more generally expressed genes, as this will reduce the efficiency of selection (Dapper and Wade 2020). However, this sort of effect requires the presence of such genes on sex chromosomes, which *Daphnia* do not have. Indeed, the fact that silent-site diversities for genes expressed in the testes, whether general or specific in expression, is very similar to that for other genes, suggests that the effective population size of such genes is comparable to that for genes expressed in female tissues.

Despite these deviations of *Daphnia* mating systems relative to those in obligately sexual species, the unique evolutionary features of *Daphnia* testes-specific genes are similar to what has been observed in other species. For example, elevated numbers of testes-specific lncRNAs have been found in *Drosophila* (Vedelek et al. 2018). Testes-specific protein-coding genes are also known to have high levels of amino-acid sequence evolution and high turnover rates in different fly lineages, albeit with much higher levels of *d*_N_ /*d*_S_ (often >1.0) than observed here (Zhang et al. 2004; Wagstaff and Begun 2005; Haerty et al. 2007; Findlay et al. 2009). Elevated rates of evolution of reproductive proteins have been noted in many other organisms, although some of this enhancement may be a result of relaxed selection (Swanson and Vacquier 2002; Turner et al. 2008; Pierron et al. 2013; Zhao et al. 2023).

All this being said, in terms of within-population diversity, whereas for protein-coding genes with *D. obtusa* orthologs, π_N_/π_S_ is inflated by 40% for the testes-specific genes relative to those restricted to other tissues, π_N_/π_S_ is not exceptional for those without *D. obtusa* orthologs (**Table 1**), and testes-specific lncRNAs have levels of π very similar to the averages for other tissue-specific genes (**Table 3**). On the other hand, *d*_N_ and *d*_N_/*d*_S_ for testes-specific genes are elevated by 50% relative to levels for genes with expression restricted to different tissues, but divergence levels for testes-specific lncRNAs are not unusual.

More generally, it is notable that much higher fractions of tissue-specific genes have no orthologs in the outgroup species than have them (approximately two thirds), whereas for genes with general expression, this fraction is just 15% (**Table 1**). This pattern is particularly prominent for testes-specific genes, where those without orthologs in *D. obtusa* are 30% more abundant than those with orthologs. Approximately 21.5% of the testis-specific genes in *D. pulex* do have orthologs in *D. obtusa*. As pools of genes lacking orthologs are likely to be enriched with young genes, this is consistent with the idea that *de novo* genes typically have restricted expression patterns (Brown et al. 2014; Kondo et al. 2017), if for no other reason than not having yet developed transcription-factor binding sites essential for broad expression. More specifically, our observations are consistent with the argument that the male germline may be permissive (or at least a filter) for the expression of novel genes, paving the way for the gradual integration into more general expression patterns (Kaessmann 2010; Rödelsperger et al. 2021; Ma et al. 2024). Such a condition might be especially true in *Daphnia*, where males constitute a very small proportion of populations. Further comparative studies across multiple *Daphnia* species will be necessary to determine the directionality of patterns of tissue specificity of gene expression.

#### Visual-system evolution

One of the more notable features of *Daphnia* is the large, black cyclopean eye, which is sensitive to alternative wavelengths and intensities of light, and is used to guide short-term behaviors such as diel vertical migration and longer-term behaviors such as diapause (Consi and Macagno 1985; Ringelberg 1999; Weiss et al. 2012; Slusarczyk and Flis 2019; Chen et al. 2024). Given the relative ease of quantifying such responses, the *Daphnia* visual system thus provides a useful model for evaluating the cellular mechanisms underlying light-triggered behavior. To this end, we examined the tissue-specific transcriptome profiles to identify genes enriched in the *Daphnia* eye. Of the 10,224 genes showing expression in the eye, 40 have eye-specific expression (**Supplementary File**), including two opsin genes, LOC124188658 and LOC124203478.

Opsins are the key light-sensitive G-protein coupled receptors used in metazoan vision, and these expanded into multiple functional groups prior to the emergence of the genus *Daphnia* and in many cases are pan-crustacean (Colbourne et al. 2011; Brandon et al. 2017; Palecanda et al. 2022). The resultant groups include multiple forms of visual opsins (sensitive to various wavelengths of light), arthropsins (whose functions remain unknown), and ciliary opsins. Based on these prior genome annotation efforts, we anticipated the presence of 46 to 48 opsins in *D. pulex*, and our analyses of the KAP4 transcriptome reveal 46 opsins distributed over the three major functional classes (**Figure 3**). Of these 46, 30 are visual opsins, including nine green wavelength-sensitive opsins (LOPA), 17 red wavelength-sensitive opsins (LOPB), one blue wavelength-sensitive opsin (BLOP), one ultraviolet wavelength-sensitive opsin (UVO), and two opsins of unknown sensitivity (UNOP) (**Figure 3**). Our expression data indicate that all 30 are expressed in the adult eye, although we do not know how these are distributed over the 22 ommatidia and 176 photoreceptors contained within the *Daphnia* eye (Macagno et al. 1973). Whereas the functions of arthropsins remain elusive, our data indicate that their expression is concentrated in the brain along with ciliary opsins (**Figure 3**), which have been implicated in circadian entrainment (Arendt et al. 2004; Velarde et al. 2005).

**Figure 3.**
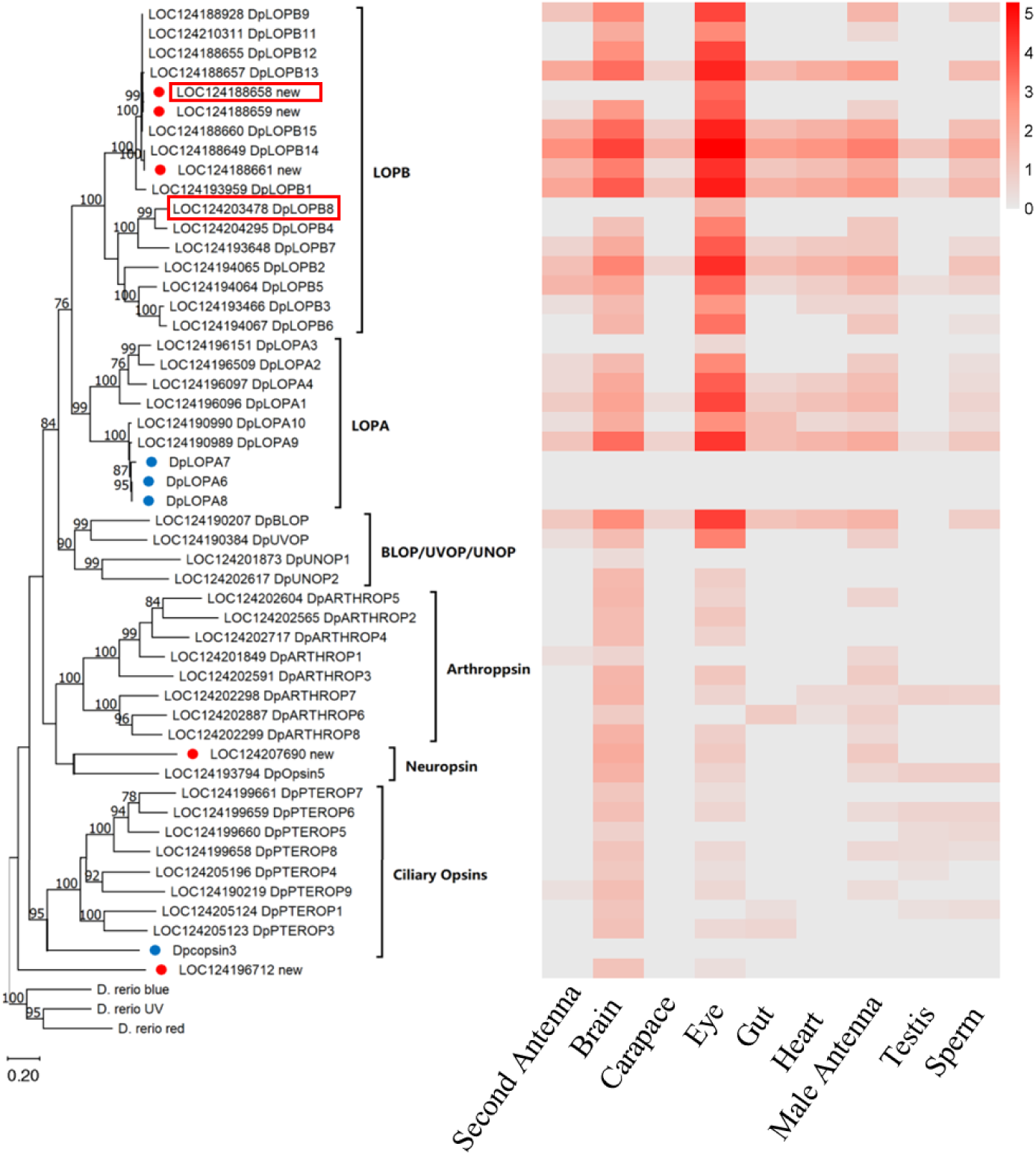
Identification and phylogenetic analysis of *Daphnia pulex* opsin genes. The maximum-likelihood tree was constructed using MEGA X (Kumar et al. 2018) with default settings, based on the full-length DNA sequences of *D. pulex* opsin genes identified in this study as well as those from Brandon et al. (2017). The first part of each sequence ID corresponds to gene accessions in the KAP4 genome assembly, while the second part represents sequences retrieved from Brandon et al. (2017). Red circles indicate opsin genes newly identified in this study, whereas the four blue circles represent opsin genes identified in Brandon et al. (2017) that are absent from the KAP4 genome assembly (their expression levels are unknown). Genes in red rectangles indicate eye-specific genes. Outgroup sequences are zebrafish blue-sensitive opsin, red-sensitive opsin, and ultraviolet-sensitive opsins. The heatmap is generated using log10^(TPM+1)^, where TPM denotes transcripts per million. Red colors indicate higher expression levels and gray indicate lower expression levels.

As mentioned above, our expression data suggest that only two of the visual opsins are eye-specific (**Figure 3**), and consistent with this, we also find that many other genes that are expected to be involved in the phototransduction cascade (based on studies in *Drosophila*; Tian et al. 2012) are enriched in the eye but not eye-specific. For example, norpA, which encodes phosphatidylinositol phospholipase and is absolutely necessary for phototransduction, and arrestin 1 and arrestin 2 (arr1 and arr2), which are required for turning off activated opsins, are expressed in all other tissues tested (**Supplementary File**). In addition, we find eight of the long-wave visual opsins (two LOPA and six LOPB) and the single blue-sensitive opsin to be expressed in the carapace. These spatial data are in agreement with single-cell transcriptome analysis of whole adult *D. magna*, which led to the suggestion that the *Daphnia* cuticle may have photoreceptor-like light-detecting properties (Krishnan et al. 2024). Altogether, the phylogenetic tree of opsins (**Figure 3**) combined with an overlay of tissue-specific expression levels (**Supplementary File**) provides insight into the cellular/developmental evolutionary assignments of genes with alternative functions.

### Expression Dynamics of Gene Duplicates

Gene duplication can be a major driver of evolutionary innovation, providing raw material for the development of new functions. Understanding how expression levels change following duplication can shed light on the mechanisms of gene retention and the roles these genes play in adaptation. To explore the evolutionary fate of recently duplicated genes, we identified gene pairs in *D. pulex* that have undergone recent duplication events, specifically those genes with two copies in *D. pulex* but only a single copy in the outgroup species *D. obtusa* and *D. magna*. A total of 74 pairs of such gene duplicates were identified. For each duplicate pair, we evaluated which copy was more closely related to the *D. obtusa* ortholog at the sequence level, and then compared the expression levels of the two copies across various tissues. On average, expression levels of the more divergent copies were found to be between 50% and 67% of those of the less divergent copies (**Figure 4A**): 57% of the 74 pairs displayed higher expression in the less divergent copy, 26% showed no significant difference between the two copies, and 17% exhibited higher expression levels in the more divergent copy (**Figure 4B; Supplementary Figure S1**).

**Figure 4.**
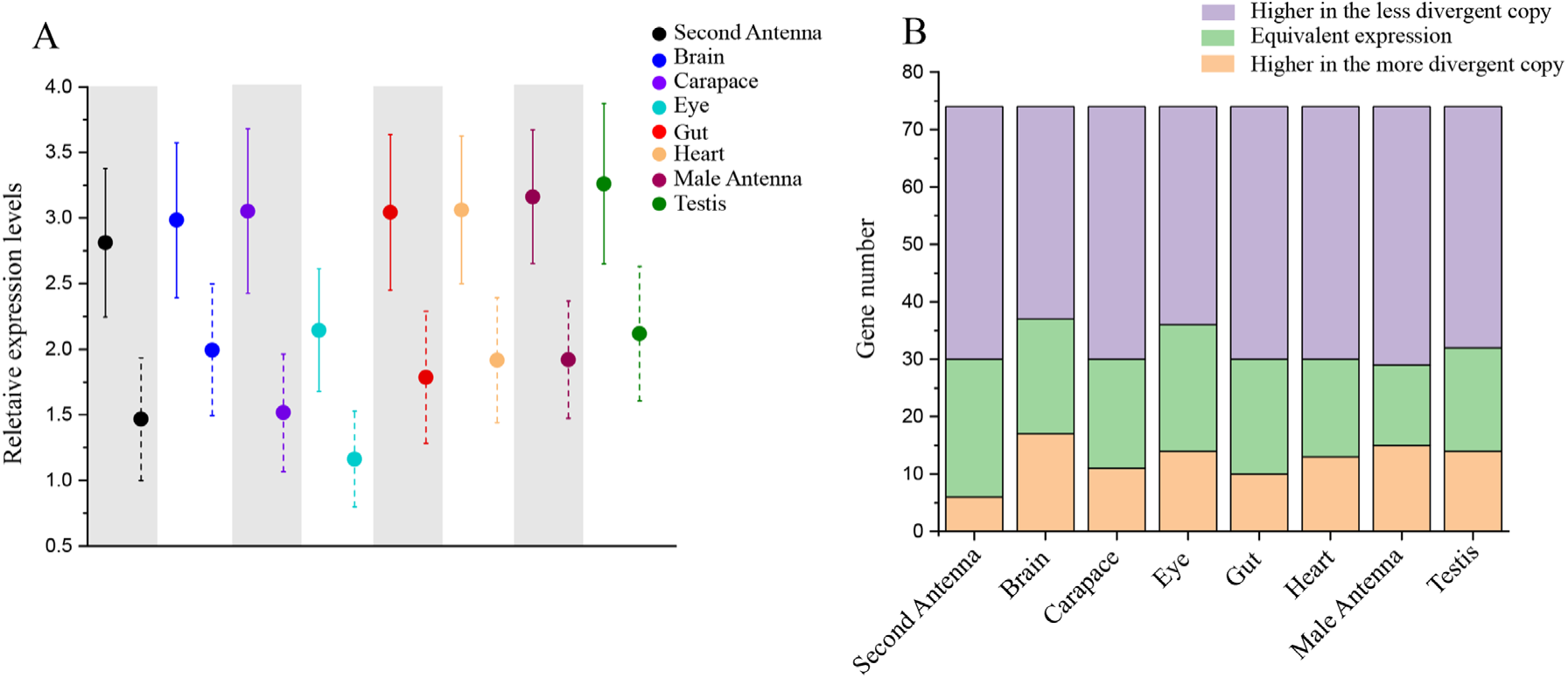
Expression levels of duplicated genes across various tissues. **A**) Average expression levels of duplicated genes: Data points are presented as mean ± 95% confidence intervals (CI). Circles with solid bars represent less divergent gene copies, while those with dashed bars represent the more divergent copies. Divergence is determined based on sequence distance from orthologs in *D. obtusa*. The y-axis displays relative expression levels in log10^(TPM+1)^. In all cases, the less divergent copy exhibits significantly higher expression levels compared to the more divergent copy (*P* < 0.05, t-test). **B**) Comparison of gene expression between less and more divergent gene copies: if the expression difference between the two copies is within 5% of the less divergent gene’s expression, they are considered to have equivalent expression levels.

These differences in expression levels might reflect functional divergence, but also could be a consequence of relaxed selection pressure on single members of the pairs. To further evaluate this issue, we examined whether the expression differences between copies are correlated with synonymous sites (*d*_S_), nonsynonymous changes (*d*_N_), and *d*_N_/*d*_S._ Our analysis revealed no significant correlation between *d*_S_ and expression-level differences (**Supplementary Figure S2**), which is not surprising if silent sites are evolving in a neutral fashion. When analyzing *d*_N,_ we observed that expression level differences in the heart and carapace tissues were significantly associated with nonsynonymous sequence divergence (**Supplementary Figure S3**), indicating that functional changes in the coding regions may contribute to tissue-specific expression divergence. Furthermore, the significant correlation between *d*_N_/*d*_S_ values and expression differences in the carapace (**Supplementary Figure S4**) suggests an alteration in the mode of selection on some gene duplicates in this tissue. We also searched for structural differences between the duplicate-gene copies, such as variations in intron-exon structures, but no substantial differences were detected. This suggests that expression divergence between duplicates is likely driven by regulatory changes rather than structural modifications. Overall, this study sheds light on the dynamics of gene duplication in *D. pulex*, highlighting that the most divergent gene copies exhibit reduced expression. Further investigations into the regulatory mechanisms underlying these changes could provide deeper insights into the evolutionary trajectories of gene duplicates.

## Discussion

This combined analysis of tissue-specific expression data with extensive population-genetic and sequence-divergence estimates provides an unprecedented view of the general nature of selection operating on genes with alternative patterns of expression in *D. pulex*. Further, it has been possible to subdivide analyses into genes: lacking orthologs in a closely related outgroup species (which are likely either novel genes in *D. pulex* and/or expendable genes lost from *D. obtusa*); genes with putative known functions (based on orthologs from model metazoan species) vs. those that remain unannotated; and protein-coding vs. lncRNA genes. These gene-specific observations provide a useful database for future studies of *Daphnia* biology, as previously accomplished in the fly *Drosophila* (Li et al. 2022) and the nematode *Caenorhabditis* (Ghaddar et al. 2023).

Several generalizations can be made with respect to protein-coding genes. First, across all tissues, protein-coding genes without orthologs in *D. obtusa* have four-fold elevations of π_N_/π_S_ (Table 1). These genes are unlikely to result from recent gene duplications, as such duplications in *D. pulex* typically retain orthologs in *D. obtusa*. Focusing on the 1,658 genes expressed in *D. pulex* that lack orthologs in *D. obtusa*, we searched for their presence in *D. pulicaria* (a sister species of *D. pulex*) and *D. magna* (a distant outgroup, in a different subgenus, to the former three). Among these genes, 944 appear to be losses from *D. obtusa*, as they are present in all three other species, while 227 are predicted to be gains in *D. pulex*, as they are unique to this species. The evolutionary history of the remaining 487 genes in this pool could not be inferred.

Second, protein-coding genes that are unannotated with respect to putative functions appear to be experiencing some form of selection based on the population data on both polymorphism and divergence, and hence are likely biologically relevant genes (as opposed to being mis-annotations). Those without *D. obtusa* orthologs have a 2.6× inflation in π_N_/π_S_, but still with a mean value of the latter (0.78) significantly below the neutral expectation of 1.0. Those with no evidence of expression in the tissues examined but nonetheless with *D. obtusa* orthologs have average π_N_/π_S_ ≃ 0.55, also well below the neutral expectation. Only a small fraction the total pool of inferred protein-coding genes (158 in total, and < 1% of the total pool of such genes) with no expression and no *D. obtusa* orthologs have average π_N_/π_S_ consistent with neutrality, and thus are candidates for pseudogene status.

Third, as a general class, tissue-specific protein-coding genes with *D. obtusa* orthologs have a 40% inflation of π_N_/π_S_ and a 70% inflation of *d*_N_ /*d*_S_. Thus, a substantial fraction of this particular class of genes is likely to be well-integrated into the biology of these species and to be experiencing positive selection for functional changes since the divergence between the two (likely at least 5 million years ago, based on silent-site divergence).

Fourth, many of the genes with *d*_N_ /*d*_S_ significantly >1.0 also have π_N_/π_S_ in excess of 1.0, suggesting selection encouraging both within-population diversity (some form of balancing selection) and between-species divergence. Here, it should be noted that these two patterns can be driven by the same forces. This is because balancing selection can elevate the rate of fixation of key mutations when the equilibrium within-population frequency deviates significantly from 0.5, as this keeps alleles from drifting to intermediate frequencies where they are otherwise protected from loss/fixation (Robertson 1962).

Fifth, for all tissues and classes of genes, there are negative correlations between expression levels and polymorphism and divergence statistics. Such observations are consistent with the common view that protein-coding genes with sequences that encourage misfolding and/or inappropriate protein-protein interactions are more deleterious when highly expressed (reviewed in Lynch 2024). It should be noted, however, that these effects are not large, with r^2^ values generally in the range of 0.03 to 0.10.

Our evolutionary analyses of lncRNAs in *D. pulex* also yield insight into this enigmatic group of expressed but noncoding DNAs, in ways that have not been pursued in other model systems. Over the past decade, it has become increasingly clear that long-noncoding (lnc) RNA genes comprise a substantial fraction of the expressed genomes of metazoans (Kapusta and Feschotte 2014; Ulitsky 2016; Mattick et al. 2023). Although the functions of most remain unknown, and many may be nonfunctional and simply reflect transcriptional noise (Ponting and Haerty 2022), those that have been assigned functions are generally involved in the regulation of various cellular and developmental processes. The human genome encodes ∼38,000 genes, 47% of which are lncRNAs, and the *Drosophila melanogaster* genome encodes ∼16,500 genes, ∼15% of which are lncRNAs (Camilleri-Robles et al. 2022).

LncRNAs comprise ∼20% of the ∼22,000 expressed genes so far identified in *D. pulex*, more than the number observed in flies, but still consistent with the proposed reduced fractions of such genes in invertebrates vs. vertebrates (Kapusta and Feschotte 2014). As in other species, the expression level of lncRNAs in *D. pulex* is reduced relative to that for protein-coding genes, on average 38% of the latter for generally expressed genes and 30% for genes with tissue-specific expression. In flies and mammals, about half of lncRNAs reside in intergenic regions and the other half overlapping protein-coding sequence in the sense or antisense directions, whereas the relative frequencies of the two types in *Daphnia* are 77% and 23%, respectively. In addition, lncRNAs have more restricted locations of expression than protein-coding genes in *D. pulex*: 56% of protein-coding genes were expressed in at least seven of the eight tissues examined, whereas only 31% of lncRNAs had similarly broad expression; in addition, 9% of protein-coding genes but 13% of lncRNAs have tissue-specific expression.

Typically, metazoan lncRNAs evolve rapidly at the DNA-sequence level, have low rates of retention of secondary structures, and have high rates of turnover among species (Kutter et al. 2012; Kapusta and Feschotte 2014; Mattick et al. 2023), consistent with the findings here. 40% of *D. pulex* lncRNA genes with general expression patterns do not have obvious orthologs in the closely related outgroup species *D. obtusa* (which is ∼11% divergent at the level of silent sites), and this increases to 71% for lncRNAs with tissue-specific expression. Notably, there is a 17-fold increase in the numbers of tissue-specific lncRNAs in testes relative to other tissues, nearly 80% of which are without orthologs in *D. obtusa*.

However, despite the relatively volatile evolution of lncRNAs, the evidence suggests that they are under some form of selection. On average and consistent across tissues, the level of within-population sequence diversity (π) is just 60% of that for silent-sites in protein-coding genes, suggesting moderate purifying selection, although not as strong as that for amino-acid sequence. On the other hand, the level of divergence from *D. obtusa* is ∼45% that for silent sites in protein-coding genes but ∼15× that for amino-acid replacement sites. Taken together, these results suggest that a substantial fraction of sites within *D. pulex* lncRNA genes are experiencing some form of purifying selection and/or strong enough positive selection to reduce within-population diversity by selective sweeps. Although there is a negative association between expression level and nucleotide diversity within the lncRNAs in essentially all tissues, this is much weaker than in the case of protein-coding genes, and the association at the level of nucleotide divergence is negligible. Managadze et al. (2011) have previously suggested a weak negative relationship of the latter kind in mammals.

## Methods

### *Daphnia* culturing and maintenance

A single adult female *D. pulex* from the reference strain KAP4 genome assembly (accessible at NCBI under accession GCF 021134715.1) was isolated from the Lynch Lab clone bank to start the culture used throughout this study. This strain was validated molecularly using unique mitochondrial genome markers, and then expanded in 250 mL beakers of a mixture of synthetic medium and sterilized lake water (from Saguaro Lake). The animals were fed daily with a pure culture of *Scenedesmus obliquu*s. To prevent overcrowding, animals were transferred weekly into new beakers. All cultures were maintained in a 20°C room under a 12:12-hour light: dark cycle.

### Tissue identification and dissection

Tissues were obtained through manual dissection, with 100 adult *Daphnia* individuals used per tissue sample, and all tissues were immediately stored in RNA Shield (Catalog No. R1100-250) to preserve RNA integrity. The dissection process involved using fine forceps to carefully grasp the *Daphnia*, cutting open the exoskeleton to expose the internal tissues. We isolated the specific tissues of interest (e.g., eye, testis, gut) while ensuring that only the target tissue was collected. Care was taken to avoid contamination from surrounding tissues. To prevent RNA degradation, the dissected tissues were immediately placed into 2 ml tubes containing RNA Shield. This step helped preserve the RNA by inhibiting RNase activity. To avoid tissue contamination, we ensured that only one tissue type was handled at a time. Each tissue was dissected separately, and the dissection bench was thoroughly cleaned before each procedure to prevent cross-contamination. The dissection protocol is documented in video format at https://www.youtube.com/@LynchLabCME. For each tissue, three replicates were performed, with each replicate generated from 100 individuals. These replicates were independently dissected, and RNA extraction and sequencing were carried out separately to ensure accurate and reproducible results.

### Total RNA isolation and sequencing

Zymo RNA purification kits were used to extract total RNA from each tissue. Prior to starting an extraction, 300 µL of RNA Secure was added to the tissue lysate (prepared in 700 µL of Zymo RNA Lysis Buffer) to help prevent RNA degradation. Final concentrations were checked on a Qubit, and RNA integrity was checked on an Agilent TapeStation. NEBNext Single Cell/Low Input RNA Library Prep Kit for Illumina was used to construct RNA-seq libraries. After cDNA synthesis and library preparation, cDNA quantities were checked using a Qubit, and samples were pooled in equimolar concentrations to ensure equal representation of each sample in the sequencing run. Each library was uniquely barcoded during preparation to enable pooling and differentiation of samples during downstream bioinformatics analyses. Sequencing was performed on an Illumina NovaSeq 6000 platform, generating paired-end reads with a read length of 150 bp. A total of approximately 50 million reads per sample were obtained, providing sufficient depth for downstream transcriptomic analysis.

### RNA-seq read mapping and tissue-specific expression

Raw reads from each sample were filtered using Trimmomatic (version 0.36) (Bolger et al. 2014) with the settings “AVGQUAL:20 ILLUMINACLIP:TruSeq3-PE.fa:2:30:10 TRAILING:20 MINLEN:50.” After trimming, reads mapped to *D. pulex* ribosomal RNA were excluded from the analysis by using the “samtools view -f 4” command. The remaining reads were mapped to the high-quality *D. pulex* genome assembly from isolate KAP4 (NCBI accession: GCF 021134715.1) using hisat2. Uniquely mapped reads were retrieved using the command “samtools view -q 60.” Relative expressions (TPM values) for each gene were obtained using StringTie (Pertea et al. 2015) with default settings. To calculate TPM, we first normalize the read count for each gene by its length in kilobases (RPK), then divide each gene’s RPK by the sum of all RPKs in the sample, and multiply by 1 million. This adjusts for both gene length and sequencing depth. To avoid spurious expression signals, protein-coding genes with TPM < 1 were considered not expressed. Tissue-specific protein-coding genes were defined as those with an average TPM > 1 in one tissue and TPM < 1 in all other tissues. For lncRNAs, a cutoff of 0.2 was applied, as these are generally expressed at lower levels.

lncRNAs were predicted by NCBI using by Eukaryotic Genome Annotation Pipeline from NCBI, using RNA sequencing data were obtained from various experimental conditions (Ye et al. 2017). The data were trimmed and mapped to the KAP4 *D. pulex* genome. Aligned reads were assembled into transcripts using StringTie (Pertea et al. 2015). The Cmsearch (Cui et al. 2016) was used to infer whether an RNA sequence has coding ability. For the predicted lncRNAs, Rfam (https://rfam.org/) were further used to determine their families, clans, motifs, and annotation.

### Identification of gene gains and looses

To infer gene gains and losses among *D. pulex*, *D. obtusa*, and *D. magna*, we utilized orthologous group data from OrthoDB v10 (Kriventseva et al., 2019). Proteins from each species were downloaded from NCBI and used to generate orthologous clusters. For each orthologous group predicted by OrthoDB, gene gains and losses were determined as follows: Genes present in *D. pulex* but absent in the other species were classified as potential gains in *D. pulex*, while those present in the other species but absent in *D. obtusa* were considered potential losses in *D. obtusa*. To minimize annotation artifacts, only genes supported by expression evidence (TPM > 1) were included in the analysis.

### Selection analysis

To examine the selective forces acting on protein-coding genes, we utilized population-genetic indices such as π_N_ (nonsynonymous polymorphism) and π_S_ (synonymous polymorphism), *d*_N_ (nonsynonymous divergence), and *d*_S_ (synonymous divergence), as well as the neutrality index (NI) (Rand and Kann 1996). These metrics were obtained from previously published datasets by Maruki et al. (2022) and Ye et al. (2023). π_N_/π_S_ ratios were used to infer the strength and direction of selection, where values significantly above 1 indicate positive selection, while values below 1 suggest purifying selection. Similarly, the *d*_N_/*d*_S_ ratios were employed to examine long-term evolutionary trends, with emphasis on identifying genes exhibiting elevated rates of nonsynonymous substitutions that may indicate adaptive evolution. To ensure robust conclusions, analyses focused on genes with high-confidence expression data (TPM > 1) to minimize the inclusion of pseudogenes or annotation artifacts.

### GO enrichment analysis for tissue-specific expression

To identify functional roles of genes with tissue-specific expression patterns, we conducted Gene Ontology (GO) enrichment analysis. The enrichment analysis was performed using a χ² test implemented via the Perl module Statistics::ChisqIndep to assess overrepresented GO terms among these genes. To address the issue of overlapping and redundant GO terms, we employed the web tool REVIGO (http://revigo.irb.hr/) to cluster related terms and retain the most representative ones. Overlapping GO terms with >80% shared genes were removed, ensuring a more interpretable and concise set of enriched terms. Statistical significance of enrichment was evaluated using the Benjamini-Hochberg method to correct for multiple testing, controlling the false discovery rate (FDR) at 5%.

### Data accession

The *D. pulex* (KAP4) assembly can be accessed at NCBI under accession GCF 021134715.1 and annotation at https://www.ncbi.nlm.nih.gov/genome/annotationeuk/Daphnia pulex/100/. All of the RNA sequencing reads generated in this study can be accessed at the NCBI Short Read Archive under study accession number PRJNA1164005.

## Acknowledgments

Support was provided by grants from the National Institutes of Health (2R35GM122566) to ML, the US National Science Foundation (DBI-2119963; ML) and (IOS-1922914; ML and ACZ), and Chinese National Natural Science Foundation Grant 32471695 to ZY.

## References

Arendt, D., Tessmar-Raible, K., Snyman, H., Dorresteijn, A. W., and Wittbrodt, J. 2004. Ciliary photoreceptors with a vertebrate-type opsin in an invertebrate brain. Science 306: 869–871.

Brandon, C. S., Greenwold, M. J., and Dudycha, J. L. 2017. Ancient and recent duplications support functional diversity of *Daphnia* opsins. J. Mol. Evol. 84: 12–28.

Brown, J. B., Boley, N., Eisman, R., May, G. E., Stoiber, M. H., Duff, M. O., Booth, B. W., et al. 2014. Diversity and dynamics of the *Drosophila* transcriptome. Nature 512: 93–399.

Camilleri-Robles, C., Amador, R., Klein, C. C., Guigó, R., Corominas, M., and Ruiz-Romero, M. 2022. Genomic and functional conservation of lncRNAs: lessons from flies. Mamm. Genome 33: 328–342.

Chen, L., Gómez, R., Horstmann, M., and Weiss, L. C. 2024. Temperature and light timing effects on diapause progression in *Daphnia magna*. Freshwater Biol. 69: 1596–1606.

Cheng, H., Concepcion, G. T., Feng, X., Zhang, H., and Li, H. 2021. Haplotype-resolved de novo assembly using phased assembly graphs with hifiasm. Nat. Methods 18: 170– 175.

Colbourne, J. K., Pfrender, M. E., Gilbert, D., Thomas, W. K., Tucker, A., Oakley, T. H., Tokishita, S., et al. 2011. The ecoresponsive genome of *Daphnia pulex*. Science 331: 555–561.

Consi, T. R., and Macagno, E. R. 1985. The spectral sensitivity of eye movements in response to light flashes in *Daphnia magna*. J. Comp. Physiol. A 156: 135–143.

Conway, J. R., Lex, A., and Gehlenborg, N. 2017. UpSetR: an R package for the visualization of intersecting sets and their properties. Bioinformatics 33: 2938–2940.

Cui, X., Lu, Z., Wang, S., Wang, J.-Y., and Gao, X. 2016. CMsearch: simultaneous exploration of protein sequence space and structure space improves not only protein homology detection but also protein structure prediction. Bioinformatics 32: i332– i340.

Dapper, A. L., and Wade, M. J. 2020. Relaxed selection and the rapid evolution of reproductive genes. Trends Genet. 36: 640–649.

Drummond, D. A., and Wilke, C. O. 2008. Mistranslation-induced protein misfolding as a dominant constraint on coding-sequence evolution. Cell 134: 341–352.

Durand, N. C., Shamim, M. S., Machol, I., Rao, S. S., Huntley, M. H., Lander, E. S., and Aiden, E. L. 2016. Juicer provides a one-click system for analyzing loop-resolution Hi-C experiments. Cell Syst. 3: 95–98.

Findlay, G. D., MacCoss, M. J., and Swanson, W. J. 2009. Proteomic discovery of previously unannotated, rapidly evolving seminal fluid genes in *Drosophila*. Genome Res. 19: 886–896.

Gabaldon, T., and Koonin, E. V. 2013. Functional and evolutionary implications of gene orthology. Nat. Rev. Genet. 14: 360–366.

Ghaddar, A., Armingol, E., Huynh, C., Gevirtzman, L., Lewis, N. E., Waterston, R., and O’Rourke, E. J. 2023. Whole-body gene expression atlas of an adult metazoan. Sci. Adv. 9: eadg0506.

Guan, D., McCarthy, S. A., Wood, J., Howe, K., Wang, Y., and Durbin, R. 2020. Identifying and removing haplotypic duplication in primary genome assemblies. Bioinformatics 36: 2896–2898.

Haerty, W., Jagadeeshan, S., Kulathinal, R. J., Wong, A., Ram, K. R., Sirot, L. K., Levesque, L., et al. 2007. Evolution in the fast lane: rapidly evolving sex-related genes in *Drosophila*. Genetics 177: 1321–1335.

Kaessmann, H. 2010. Origins, evolution, and phenotypic impact of new genes. Genome Res. 20: 1313–1326.

Kaletsky, R., Yao, V., Williams, A., Runnels, A. M., Tadych, A., Zhou, S., Troyanskaya, O. G., and Murphy, C. T. 2018. Transcriptome analysis of adult *Caenorhabditis elegans* cells reveals tissue-specific gene and isoform expression. PLoS Genet. 14: e1007559.

Kapusta, A., and C. Feschotte. 2014. Volatile evolution of long noncoding RNA repertoires: mechanisms and biological implications. Trends Genet. 30: 439–452.

Klein, M., B. Afonso, A. J. Vonner, L. Hernandez-Nunez, M. Berck, C. J. Tabone, E. A. Kane, V. A. Pieribone, M. N. Nitabach, et al. 2015. Sensory determinants of behavioral dynamics in *Drosophila* thermotaxis. Proc. Natl. Acad. Sci. USA 112: E220–E229.

Kondo, S., J. Vedanayagam, J. Mohammed, S. Eizadshenass, L. Kan, N. Pang, R. Aradhya, A. Siepel, J. Steinhauer, and E. C. Lai. 2017. New genes often acquire male-specific functions but rarely become essential in *Drosophila*. Genes Dev. 31: 1841–1846.

Krishnan, I., L. Y. Yampolsky, K. Petrova, and L. Peshkin. 2024. Single-cell transcriptome defines cell type repertoire of adult *Daphnia magna*. bioRxiv. 10.1101/2024-05.

Kriventseva EV, Kuznetsov D, Tegenfeldt F, Manni M, Dias R, Simão FA, Zdobnov EM. 2019. OrthoDB v10: sampling the diversity of animal, plant, fungal, protist, bacterial, and viral genomes for evolutionary and functional annotations of orthologs. Nucleic Acids Res. 47:D807–D811.

Kumar, S., G. Stecher, M. Li, C. Knyaz, and K. Tamura. 2018. MEGA X: molecular evolutionary genetics analysis across computing platforms. Mol. Biol. Evol. 35: 1547– 1549.

Kutter, C., S. Watt, K. Stefflova, M. D. Wilson, A. Goncalves, C. P. Ponting, D. T. Odom, and A. C. Marques. 2012. Rapid turnover of long noncoding RNAs and the evolution of gene expression. PLoS Genet. 8: e1002841.

Lex, A., N. Gehlenborg, H. Strobelt, R. Vuillemot, and H. Pfister. 2014. UpSet: visualization of intersecting sets. IEEE Trans. Vis. Comput. Graph. 20: 1983–1992.

Li, H., J. Janssens, M. De Waegeneer, S. S. Kolluru, K. Davie, V. Gardeux, W. Saelens, F. P. David, M. Brbic, K. Spanier, et al. 2022. Fly Cell Atlas: a single-nucleus transcriptomic atlas of the adult fruit fly. Science 375: eabk2432.

Lynch, M. 2007. The Origins of Genome Architecture. Sinauer Associates, Sunderland, MA.

Lynch, M. 2024. Evolutionary Cell Biology: The Origins of Cellular Architecture. Oxford University Press, Oxford, UK.

Lynch, M., W. Wei, Z. Ye, and M. Pfrender. 2024. The genome-wide signature of short-term temporal selection. Proc. Natl. Acad. Sci. USA 121: e2307107121.

Ma, F., C. Y. Lau, and C. Zheng. 2024. Young duplicate genes show developmental stage- and cell type-specific expression and function in *Caenorhabditis elegans*. Cell Genomics 4: 1.

Macagno, E. R., V. Lopresti, and C. Levinthal. 1973. Structure and development of neuronal connections in isogenic organisms: variations and similarities in the optic system of *Daphnia magna*. Proc. Natl. Acad. Sci. USA 70: 57–61.

Managadze, D., I. B. Rogozin, D. Chernikova, S. A. Shabalina, and E. V. Koonin. 2011. Negative correlation between expression level and evolutionary rate of long intergenic noncoding RNAs. Genome Biol. Evol. 3: 1390–1404.

Maruki, T., Z. Ye, and M. Lynch. 2022. The population genomics of a subdivided species. Mol. Biol. Evol. 9: msac152.

Mattick, J. S., P. P. Amaral, P. Carninci, S. Carpenter, H. Y. Chang, L. L. Chen, R. Chen, C. Dean, M. E. Dinger, K. A. Fitzgerald, et al. 2023. Long non-coding RNAs: definitions, functions, challenges and recommendations. Nat. Rev. Mol. Cell Biol. 24: 430–447.

Nystrom, E. E. L., L. Arike, E. Ehrencrona, G. C. Hansson, and M. E. V. Johansson. 2019. Calcium-activated chloride channel regulator 1 (CLCA1) forms non-covalent oligomers in colonic mucus and has mucin 2-processing properties. J. Biol. Chem. 294: 17075–17089.

Palecanda S, Iwanicki T, Steck M, Porter ML. 2022. Crustacean conundrums: a review of opsin diversity and evolution. Philos Trans R Soc B. 377:20210289.

Pertea M, Pertea GM, Antonescu CM, Chang T-C, Mendell JT, Salzberg SL. 2015. StringTie enables improved reconstruction of a transcriptome from RNA-seq reads. Nat Biotechnol. 33:290–295.

Pierron D, Razafindrazaka H, Rocher C, Letellier T, Grossman LI. 2014. Human testis-specific genes are under relaxed negative selection. Mol Genet Genomics. 289:37–45.

Ponting CP, Haerty W. 2022. Genome-wide analysis of human long noncoding RNAs: a provocative review. Annu Rev Genomics Hum Genet. 23:153–172.

Ringelberg J. 1999. The photobehaviour of Daphnia spp. as a model to explain diel vertical migration in zooplankton. Biol Rev. 74:397–423.

Robertson A. 1962. Selection for heterozygotes in small populations. Genetics. 47:1291– 1300.

Rödelßperger C, Ebbing A, Sharma DR, Okumura M, Sommer RJ, Korswagen HC. 2021. Spatial transcriptomics of nematodes identifies sperm cells as a source of genomic novelty and rapid evolution. Mol Biol Evol. 38:229–243.

Senthilan PR, Piepenbrock D, Ovezmyradov G, Nadrowski B, Bechstedt S, Pauls S, Winkler M, Möbius W, Howard J, Göpfert MC. 2012. Drosophila auditory organ genes and genetic hearing defects. Cell. 150:1042–1054.

Shen WL, Kwon Y, Adegbola AA, Luo J, Chess A, Montell C. 2011. Function of rhodopsin in temperature discrimination in Drosophila. Science. 331:1333–1336.

Ślusarczyk M, Flis S. 2019. Light quantity, not photoperiod terminates diapause in the crustacean Daphnia. Limnol Oceanogr. 64:124–130.

Sokabe T, Chen H-C, Luo J, Montell C. 2016. A switch in thermal preference in Drosophila larvae depends on multiple rhodopsins. Cell Rep. 17:336–344.

Swanson WJ, Vacquier VD. 2002. The rapid evolution of reproductive proteins. Nat Rev Genet. 3:137–144.

Tian Y, Hu W, Tong H, Han J. 2012. Phototransduction in Drosophila. Sci China Life Sci. 55:27–34.

Turner LM, Chuong EB, Hoekstra HE. 2008. Comparative analysis of testis protein evolution in rodents. Genetics. 179:2075–2089.

Ulitsky I. 2016. Evolution to the rescue: using comparative genomics to understand long non-coding RNAs. Nat Rev Genet. 17:601–614.

Velarde RA, Sauer CD, Walden KKO, Fahrbach SE, Robertson HM. 2005. Pteropsin: a vertebrate-like non-visual opsin expressed in the honey bee brain. Insect Biochem Mol Biol. 35:1367–1377.

Wagstaff BJ, Begun DJ. 2005. Molecular population genetics of accessory gland protein genes and testis-expressed genes in Drosophila mojavensis and D. arizonae. Genetics. 171:1083–1101.

Weisman CM, Murray AW, Eddy SR. 2022. Mixing genome annotation methods in a comparative analysis inflates the apparent number of lineage-specific genes. Curr Biol. 32:2632–2639.

Weiss LC, Tollrian R, Herbert Z, Laforsch C. 2012. Morphology of the Daphnia nervous system: a comparative study on Daphnia pulex, Daphnia lumholtzi, and Daphnia longicephala. J Morphol. 273:1392–1405.

Ye Z, Xu S, Spitze K, Asselman J, Jiang X, Ackerman MS, Lopez J, Harker B, Raborn RT, Pfrender ME, Lynch M. 2017. A new reference genome assembly for the microcrustacean Daphnia pulex. G3. 7:1405–1416.

Ye Z, Wei W, Pfrender M, Lynch M. 2023. Evolutionary insights from a large-scale survey of population-genomic variation. Mol Biol Evol. 40:msad233.

Zhang J, Yang JR. 2015. Determinants of the rate of protein sequence evolution. Nat Rev Genet. 16:409–420.

Zhang Z, Hambuch TM, Parsch J. 2004. Molecular evolution of sex-biased genes in Drosophila. Mol Biol Evol. 21:2130–2139.

Zhao L, Zhou W, He J, Li DZ, Li HT. 2024. Positive selection and relaxed purifying selection contribute to rapid evolution of male-biased genes in a dioecious flowering plant. eLife. 12:RP89941.

