## Supplementary Tables and Figures for "Evolutionary Analysis of Gene-expression Localization in the Model Crustacean, *Daphnia pulex*"

**Table S1.** Uniquely mapped reads for each sample. Coverage was calculated by dividing the summed length of all uniquely mapped reads by the length of the expressed regions in the genome.

| Sample | Reads | Coverage (×) |
| --- | --- | --- |
| Antennae1 | 41,697,804 | 249 |
| Antennae2 | 184,296,148 | 1,099 |
| Antennae3 | 16,609,935 | 99 |
| Brain1 | 9,743,302 | 58 |
| Brain2 | 14,090,818 | 84 |
| Brain3 | 123,647,719 | 738 |
| Carapase1 | 11,053,861 | 66 |
| Carapase2 | 9,500,263 | 57 |
| Carapase3 | 13,619,438 | 81 |
| Eye1 | 19,334,908 | 115 |
| Eye2 | 36,195,314 | 216 |
| Eye3 | 9,934,368 | 59 |
| Gut1 | 11,166,355 | 67 |
| Gut2 | 21,079,226 | 126 |
| Gut3 | 13,727,361 | 82 |
| Heart1 | 18,000,960 | 107 |
| Heart2 | 11,157,352 | 67 |
| Heart3 | 28,371,086 | 169 |
| Male_Antenula1 | 8,066,258 | 48 |
| Male_Antenula2 | 9,908,800 | 59 |
| Male_Antenula3 | 8,228,288 | 49 |
| Sperm1 | 19,763,434 | 118 |
| Sperm2 | 10,843,349 | 65 |
| Sperm3 | 11,945,509 | 71 |
| Teste1 | 9,848,156 | 59 |
| Teste2 | 18,332,987 | 109 |
| Teste3 | 16,495,132 | 98 |
| **mean** | **26,172,523** | **156** |

**Table S2.** Population genomic statistics for the 4,085 unannotated genes. Expressed indicate genes that are expressed in some tissues and unexpressed indicate genes that are not expressed in any tissue (TPM<1). With popgen data means genes have population genomics data available from Maruki et al. 2022, Ye et al. 2023, and Lynch et al. 2024. NA means data not available.

| Expression | Population genomics data | *D. obtusa* ortholog | π_N_ /π_S_ | SE(π_N_ /π_S_) | *d*_N_ /*d*_S_ | SE(*d*_N_ /*d*_S_) |
| --- | --- | --- | --- | --- | --- | --- |
| Expressed (3319) | With popgen data (2835) | With ortholog (2385) | 0.304 | 0.057 | 0.231 | 0.004 |
|  |  | Without ortholog (450) | 0.776 | 0.122 | NA | NA |
|  | No popgen data (480) | With ortholog (14) | NA | NA | NA | NA |
|  |  | Without ortholog (466) | NA | NA | NA | NA |
| Unexpressed (766) | With popgen data (426) | With ortholog (268) | 0.549 | 0.057 | 0.251 | 0.014 |
|  |  | Without ortholog (158) | 1.054 | 0.121 | NA | NA |
|  | No popgen data (340) | With ortholog (0) | NA | NA | NA | NA |
|  |  | Without ortholog (340) | NA | NA | NA | NA |

**Table S3.** 24 genes with *d*_N_ /*d*_S_ significantly greater than 1.0.

| Gene id | π_N_ /π_S_ | SE  (π_N_ /π_S_) | *d*_N_ /*d*_S_ | SE  (*d*_N_ /*d*_S_) | Description |
| --- | --- | --- | --- | --- | --- |
| LOC124194853 | 2.26 | 0.29 | 4.99 | 0.39 | DNA-directed RNA polymerase II subunit RPB1 |
| LOC124209528 | 1.23 | 0.04 | 2.36 | 0.07 | sucrase-isomaltase, intestinal-like isoform X2 |
| LOC124209891 | 0.86 | 0.09 | 1.95 | 0.13 | uncharacterized protein |
| LOC124195389 | 8.41 | 1.39 | 1.83 | 0.07 | DNA-directed RNA polymerase III subunit RPC7-like isoform X1 |
| LOC124207880 | 1.06 | 0.35 | 1.75 | 0.11 | uncharacterized protein |
| LOC124209772 | 1.74 | 0.18 | 1.56 | 0.01 | zinc finger protein 652-like isoform X1 |
| LOC124202778 | 0.85 | 0.05 | 1.55 | 0.05 | uncharacterized protein |
| LOC124201646 | 4.98 | 1.09 | 1.48 | 0.08 | uncharacterized protein |
| LOC124209511 | 2.89 | 0.28 | 1.47 | 0.02 | NADH-ubiquinone oxidoreductase 75 kDa subunit, mitochondrial-like |
| LOC124204253 | 0.60 | 0.19 | 1.44 | 0.05 | uncharacterized protein |
| LOC124209693 | 1.82 | 0.21 | 1.44 | 0.04 | transducin-like enhancer protein 4 isoform X4 |
| LOC124207310 | 2.17 | 0.45 | 1.43 | 0.08 | uncharacterized protein |
| LOC124209916 | 0.88 | 0.17 | 1.40 | 0.07 | leishmanolysin-like peptidase |
| LOC124210095 | 2.30 | 0.09 | 1.38 | 0.05 | bumetanide-sensitive sodium-(potassium)-chloride cotransporter-like |
| LOC124209510 | 0.99 | 0.18 | 1.36 | 0.05 | uncharacterized protein |
| LOC124200782 | 1.07 | 0.16 | 1.35 | 0.05 | F-box and WD repeat domain-containing 11-A-like |
| LOC124198817 | 0.54 | 0.10 | 1.30 | 0.13 | uncharacterized protein |
| LOC124208929 | 1.56 | 0.22 | 1.25 | 0.02 | calcium-activated chloride channel regulator 4A-like |
| LOC124209698 | 2.64 | 0.51 | 1.23 | 0.06 | protein groucho-like |
| LOC124197711 | 0.66 | 0.09 | 1.20 | 0.08 | uncharacterized protein |
| LOC124208808 | 3.71 | 1.43 | 1.16 | 0.06 | uncharacterized protein |
| LOC124204969 | 0.53 | 0.02 | 1.12 | 0.05 | uncharacterized protein |
| LOC124209813 | 1.15 | 0.08 | 1.08 | 0.04 | probable ATP-dependent RNA helicase DDX46 |
| LOC124209928 | 1.01 | 0.18 | 1.08 | 0.02 | uncharacterized protein |

**Figure S1.** Expression levels for the 74 duplicates genes in various tissues. The gene copy more closely related to *D. obtusa* is designated as the ancestral copy, while the other is classified as the derived copy. The Y-axis displays relative expression levels in log₂^(TPM + 1)^.

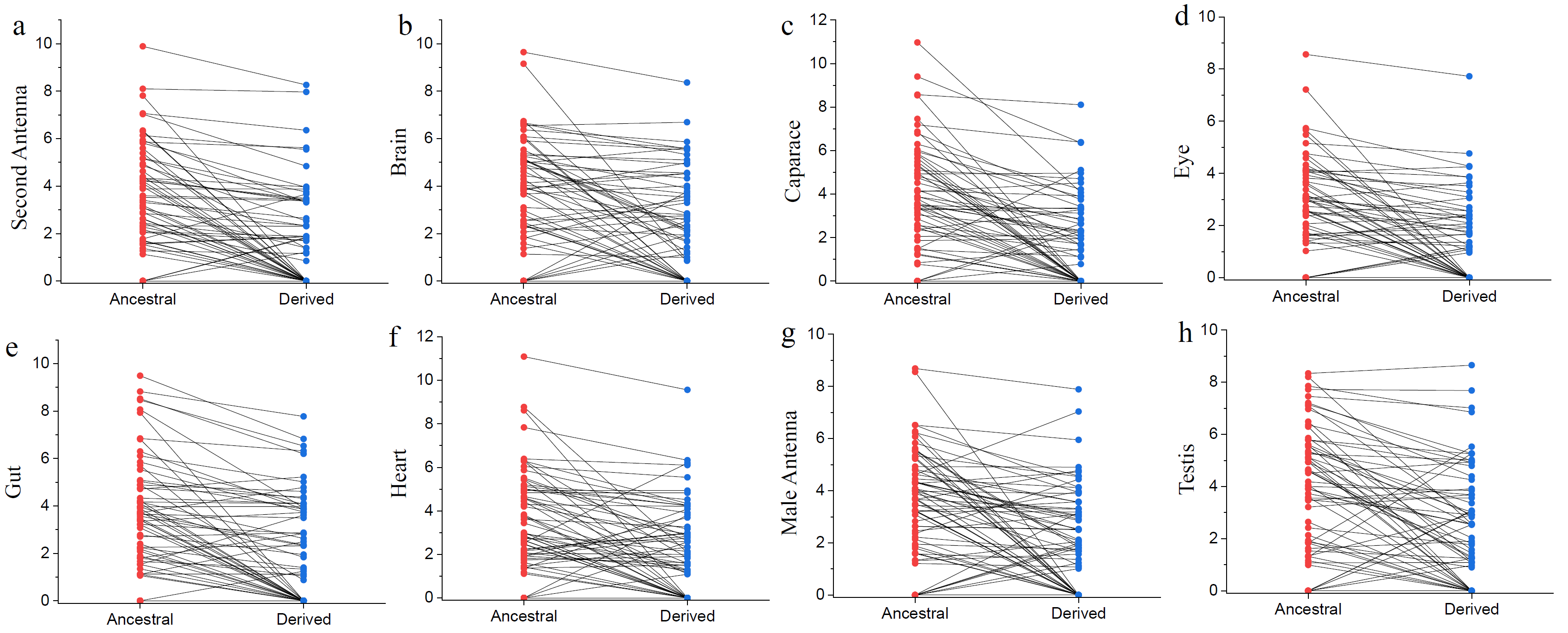

**Figure S2.** Pearson correlations between expression differences and synonymous changes (*d*_S_) in the duplicated gene copies. The X-axis represents the synonymous changes between the two gene copies, transformed using log10, while the Y-axis indicates the expression differences between them (measured in log10^TPM^).

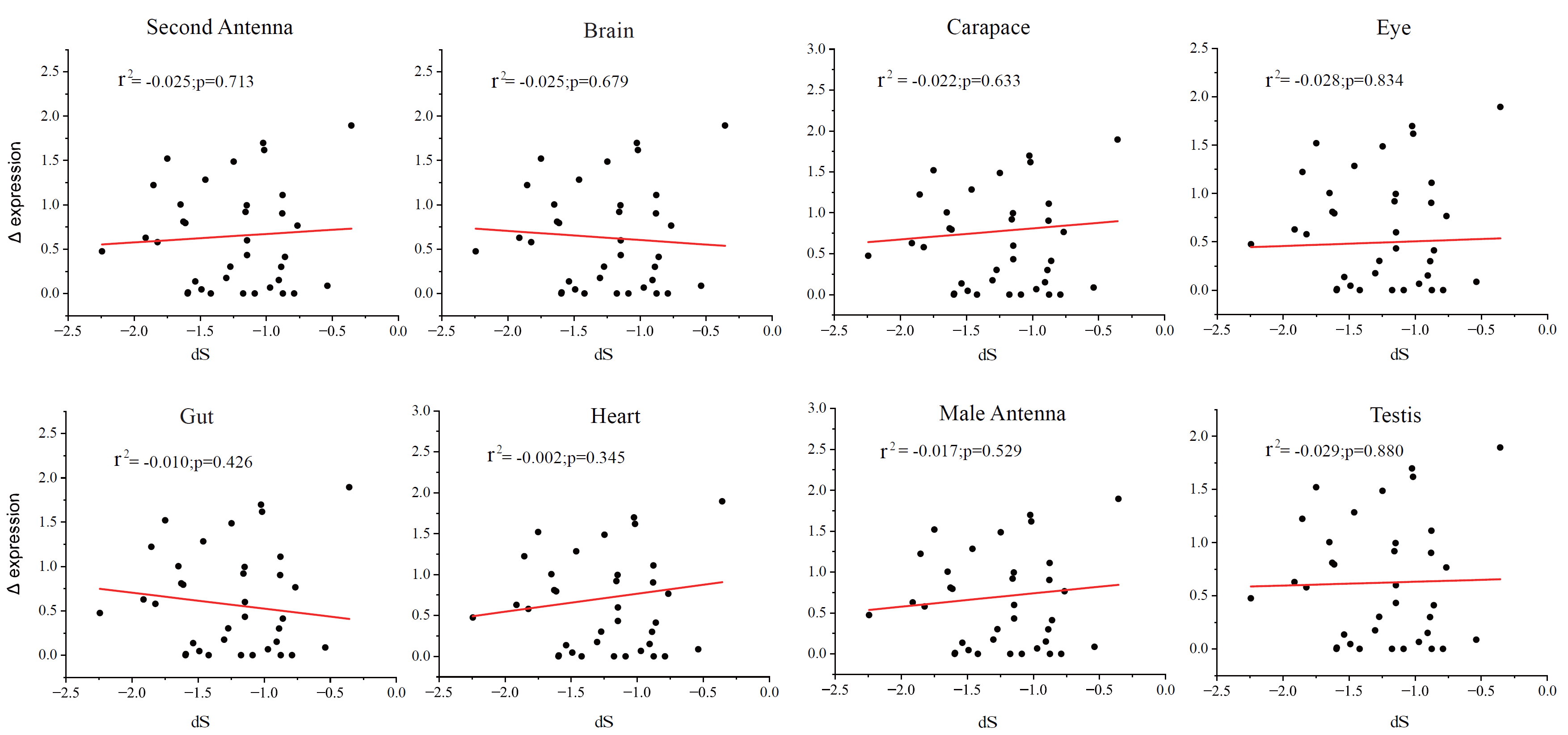

**Figure S3.** Pearson correlations between expression differences and nonsynonymous changes (*d*_N_) in the duplicated gene copies. The X-axis represents the synonymous changes between the two gene copies, transformed using log10, while the Y-axis indicates the expression differences between them (measured in log10^TPM^).

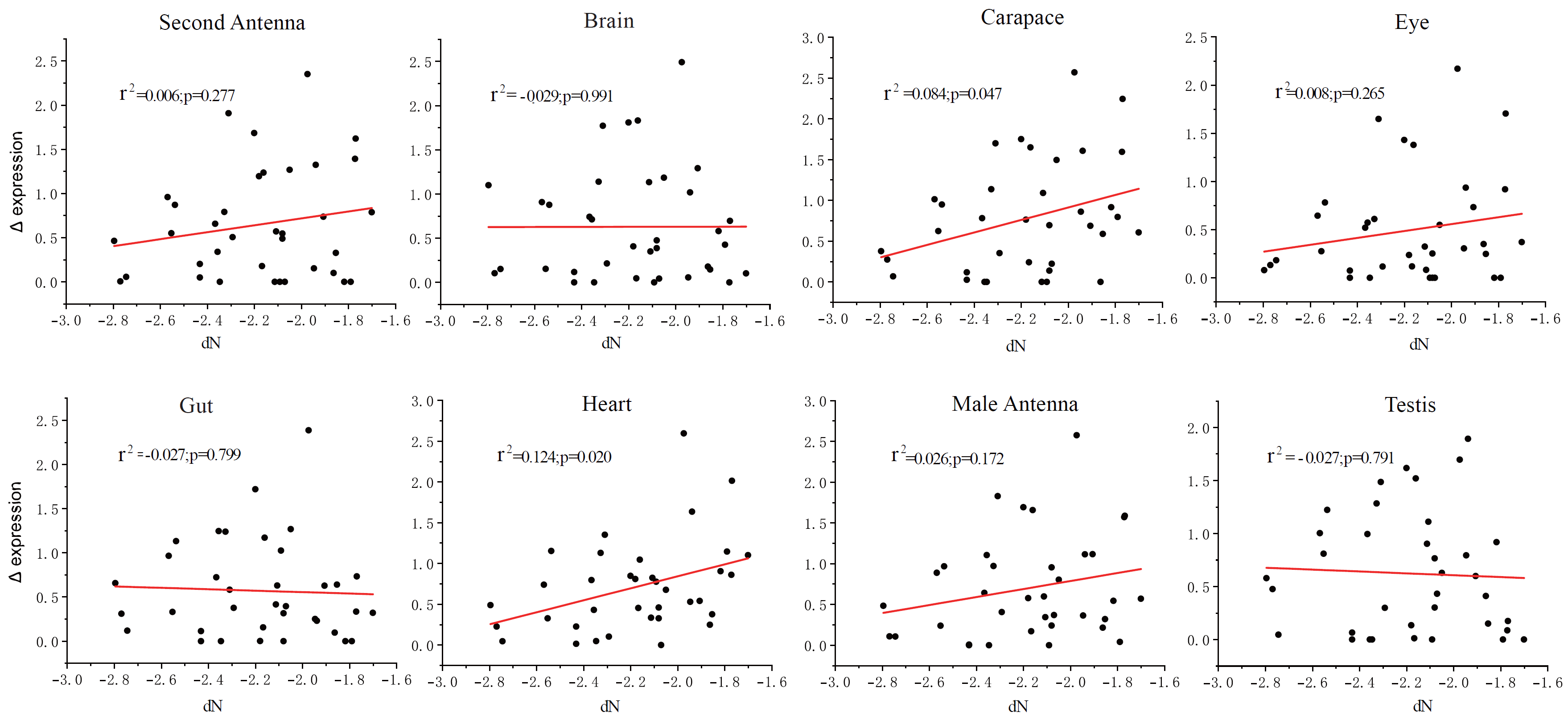

**Figure S4.** Pearson correlations between expression differences and divergence (*d*_N_/*d*_S_) in the duplicated gene copies. The X-axis represents *d*_N_/*d*_S_ differences between the two gene copies, transformed using log10, while the Y-axis indicates the expression differences between them (measured in log10^TPM^).

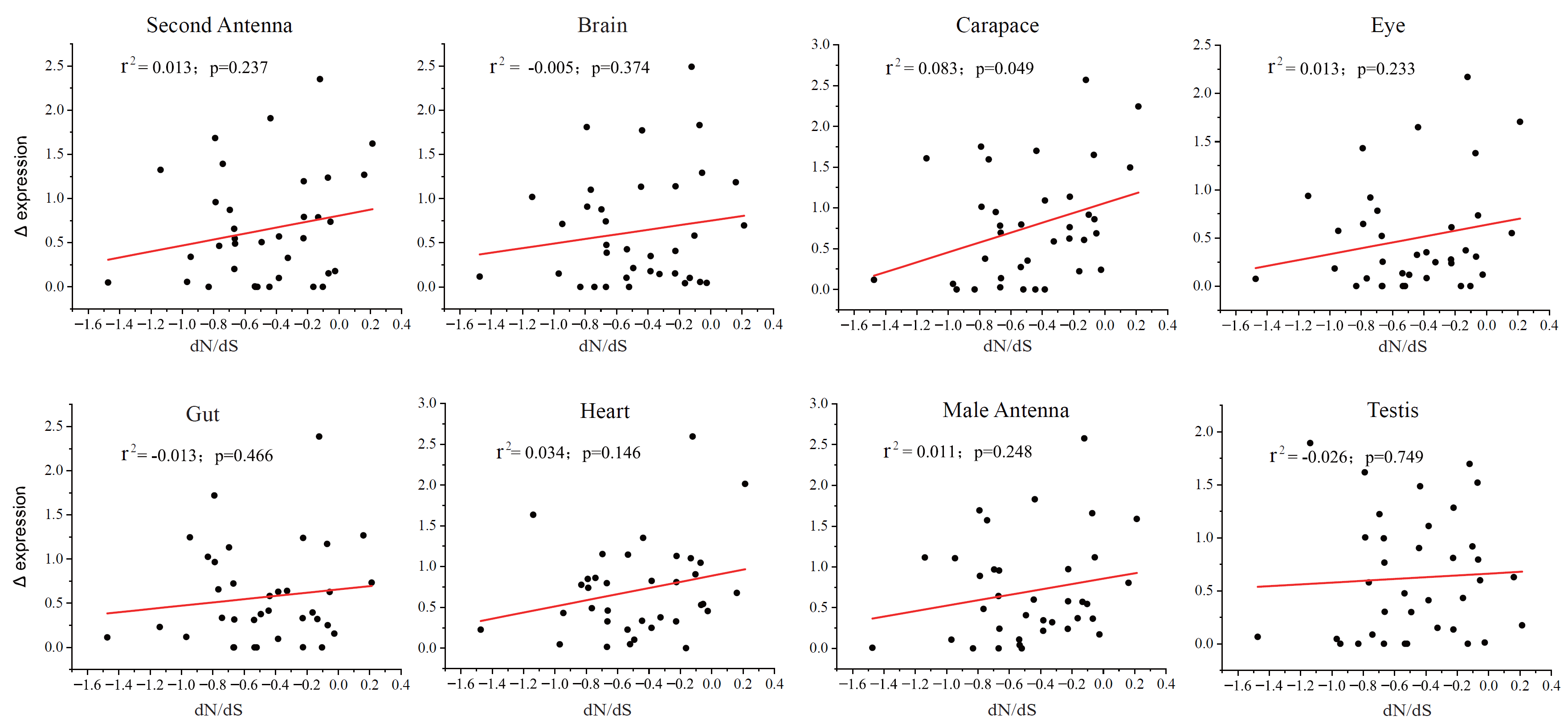

Kaletsky, R., W. Lakhina, N. Arey, S. Williams, N. Landis, A. Ashraf, J. Murphy, and C. T. Murphy. 2018. Transcriptome analysis of adult *Caenorhabditis elegans* cells reveals tissue-specific gene and isoform expression. *PLoS Genet.* 14: e1007559.

Dobson, A. J., M. E. He, C. Li, L. B. Perrone, S. M. Foster, and L. Partridge. 2018. Tissue-specific transcriptome profiling of *Drosophila* reveals roles for GATA transcription factors in longevity by dietary restriction. *NPJ Aging Mech. Dis.* 4: 5.
